# The LTB_4_-BLT1 axis regulates actomyosin and β2 integrin dynamics during neutrophil extravasation

**DOI:** 10.1101/804914

**Authors:** Bhagawat C. Subramanian, Nicolas Melis, Desu Chen, Weiye Wang, Devorah Gallardo, Roberto Weigert, Carole A. Parent

## Abstract

The eicosanoid Leukotriene B_4_ (LTB_4_) relays chemotactic signals to direct neutrophil migration to inflamed sites through its receptor BLT1. However, the mechanisms by which the LTB_4_-BLT1 axis relays chemotactic signals during intravascular neutrophil response to inflammation remain unclear. Here, we report that LTB_4_ produced by neutrophils acts as an autocrine/paracrine signal to direct the vascular recruitment, arrest and extravasation of neutrophils in a sterile inflammation model in the mouse footpad. Using Intravital Subcellular Microscopy (ISMic), we reveal that LTB_4_ elicits sustained cell polarization and adhesion responses during neutrophil arrest *in vivo*. Specifically, LTB_4_ signaling coordinates the dynamic redistribution of non-muscle Myosin IIA (NMIIA) and β_2_-integrin (Itgb2), which facilitate neutrophil arrest and extravasation. Notably, we also found that neutrophils shed extracellular vesicles (EVs) in the vascular lumen, and that inhibition of EV release blocks LTB_4_-mediated autocrine/paracrine signaling required for neutrophil arrest and extravasation. Overall, we uncover a novel complementary mechanism by which LTB_4_ relays extravasation signals in neutrophils during early inflammation response.

**SUMMARY:** Neutrophils arrest and extravasate from the blood vessels in response to infection and injury. Using intravital subcellular microscopy, Subramanian *et al*. identify a role for extracellular vesicles-based autocrine/paracrine LTB_4_-BLT1 signaling in promoting the re-arrangement of actomyosin cytoskeleton and β_2_-integrin during neutrophil extravasation in live animals.

## INTRODUCTION

Neutrophils are the first mediators of innate immune responses (Lämmermann and Kastenmüller, 2019; Liew and Kubes, 2019). As they begin their journey to reach sites of inflammation, neutrophils in blood vessels respond by rolling on and adhering to the endothelium, before extravasating to the interstitium (a process known as transendothelial migration) (Lämmermann and Kastenmüller, 2019; Zarbock and Ley, 2008), where they further migrate directionally by sensing chemotactic gradients. Neutrophils respond to primary chemoattractants, such as formylated peptides and Complement 5a (C5a), by producing the lipid mediator LTB_4_ through the activity of the 5-lipoxygenase Alox5 (Majumdar et al., 2014; Werz, 2002). LTB_4_ acts on its cognate receptor BLT1 to relay chemotactic signals to neighboring neutrophils, thereby broadening their recruitment range to the inflamed sites (Afonso et al., 2012; Lämmermann et al., 2013), where neutrophils engage with the microenvironment, secrete a variety of effector molecules, and help recruit other immune cells, to promote host defense in a timely manner (Liew and Kubes, 2019).

Studies in mice that do not produce LTB_4_ (*Alox5*^*-/-*^) or lack its receptor (*Blt1*^*-/-*^) have highlighted a critical role for the LTB_4_-BLT1 axis in mobilizing neutrophils and other myeloid cells to sites of inflammation across multiple animal models (Collin et al., 2004; Tager et al., 2000). In fact, neutrophils self-orchestrate their recruitment during arthritis using a complex chemoattractant cascade that involves LTB_4_ production in response to C5a, which proposedly functions in an autocrine manner to promote neutrophil extravasation (Miyabe et al., 2017; Sadik et al., 2012). LTB_4_ production in neutrophils has also been shown to act in a paracrine fashion for the recruitment of both neutrophils and T cells in a chronic skin allergy model in mice (Oyoshi et al., 2012). However, the precise mechanism by which the autocrine/paracrine LTB_4_-BLT1 axis functions to relay chemotactic signals leading to neutrophil extravasation has yet to be determined. To this end, we used an *in vivo* model system based on the use of ISMic, which allows the imaging of dynamic biological processes in live animals at subcellular resolution (Weigert et al., 2013; Ebrahim and Weigert, 2019). Here, we report a crucial role for neutrophil-derived LTB_4_ in regulating the dynamics of the key regulators NMIIA and Itgb2, and that EVs from neutrophils mediate the autocrine/paracrine LTB_4_-BLT1 signaling to promote neutrophil arrest and extravasation in live mice, thus complementing previously proposed mechanisms.

## RESULTS AND DISCUSSION

### The LTB_4_-BLT1 axis is required for the persistent recruitment, arrest and extravasation of neutrophils

To address the role of the LTB_4_-BLT1 axis during neutrophil extravasation, we developed an *in vivo* inflammation model by injecting heat-killed *E. coli* bioparticles (*E. coli* BPs) into the hind footpad of anesthetized mice (Fig.S1A). First, we optimized conditions to image neutrophil extravasation behavior in a mouse strain that expresses GFP in myeloid cells (*LyzM-GFP*) (Faust et al., 2000). Upon injection of *E. coli* BPs, neutrophils accumulated in the blood vessels and extravasated, whereas upon saline injection they freely flowed in the vasculature (Fig.S1A and Movie S1). A similar effect was observed using an adoptive transfer model, where neutrophils purified from the bone marrow of a *WT* mouse were labeled with a cell-permeant fluorescent dye and transferred to a recipient *WT* mouse (Fig.1A and Movie S2). To determine whether LTB_4_ generated by neutrophils is required to drive their extravasation, we inflamed the footpad of *Alox5*^*-/-*^ mice, which are incapable of producing LTB_4_, and introduced neutrophils purified from either *WT* or *Alox5*^*-/-*^ mice. We observed that up to 3.5 hours post-injection, the recruitment of *Alox5*^*-/-*^ neutrophils to the infected footpad was significantly reduced in comparison to *WT* neutrophils (Fig.1, B and C). To gain further insights into this process, we used the same model to visualize neutrophil intravascular dynamics (Movie S3). Quantitative analysis revealed that ∼60% of *WT* neutrophils displayed rolling, ∼40% showed arrest in the lumen of the blood vessels and ∼20% extravasated in response to inflammation (Fig.1, D-G). Conversely, *Alox5*^*-/-*^ neutrophils displayed significantly reduced arrest (<10%) and extravasation response (<5%), which correlated with an increased rolling response (>80%) (Fig.1, D-G). A similar defect was also observed in *Blt1*^*-/-*^ neutrophils, which lack the receptor for LTB_4_ (Fig.1, D-G). We complemented these results by investigating the intravascular dynamics of *WT, Alox5*^*-/-*^ and *Blt1*^*-/-*^ neutrophils introduced into *Blt1*^*-/-*^ mice (Movie S4 and Fig.1, H-K). As expected, *Blt1*^*-/-*^ neutrophils exhibited enhanced rolling and a significant defect in arrest and extravasation (Fig.1, H-K). On the other hand, *Alox5*^*-/-*^ neutrophils displayed rolling, arrest and extravasation responses comparable to those of *WT* neutrophils, indicating that *Alox5*^*-/-*^ neutrophils likely utilize LTB_4_ generated by resident cells, including neutrophils, to undergo extravasation in *Blt1*^*-/-*^ mice (Fig.1, H-K). This observation also suggests that under physiological conditions the tissue environment is capable of producing the LTB_4_ required for neutrophil extravasation. Collectively, these findings establish that production and sensing of LTB_4_ in neutrophils is critical for arrest and extravasation responses *in vivo*.

**Figure 1.**
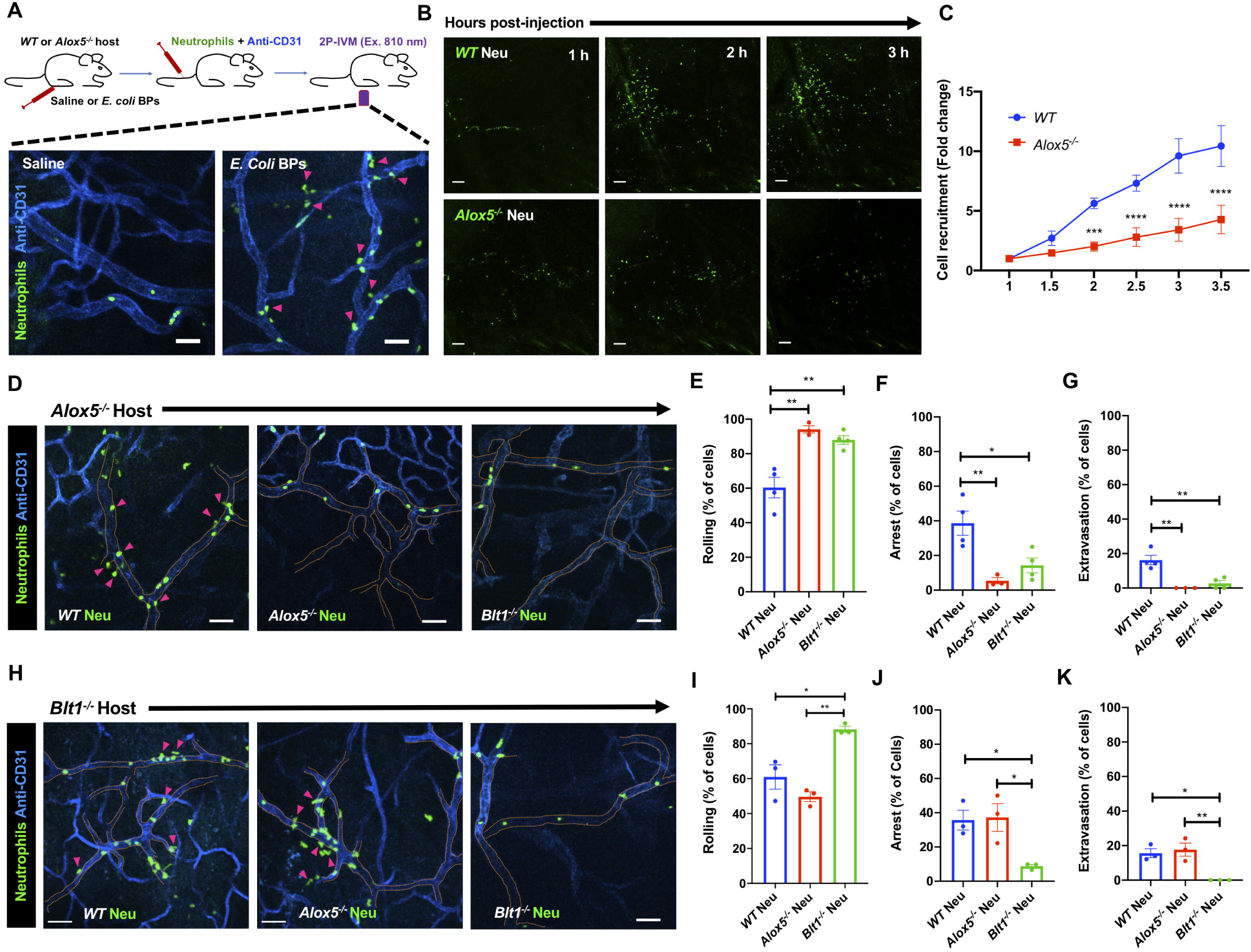
Production and sensing of LTB_4_ via BLT1 in neutrophil promotes recruitment, arrest and extravasation response *in vivo*. **(A)** Upper Panel – Diagram of the 2P-IVM procedures in the footpad of an adoptive transfer mouse model. Lower panels -Neutrophils purified from *WT* mouse strains were labeled with CellTracker Green CMFDA and adoptively transferred to recipient *WT* mice that were injected in the footpad with either saline (left panel) or *E. coli* BPs (right panel) and imaged by 2P-IVM. Maximum projections of Z-stacks are shown (vasculature in blue and neutrophils, in green). Pink arrowheads mark extravasating neutrophils (right panel). Scale bar equals 50 µm. The experiments were performed in N=2 mice. See Movie S2. **(B-C)** Neutrophils purified from either *WT* (upper panels) or *Alox5*^*-/-*^ (lower panels) mice were labeled with CellTracker Green CMFDA and adoptively transferred to *Alox5*^*-/-*^ mice with inflamed footpad. **B**-Maximum projections of Z-stacks of the footpads are presented for 1 h, 2 h and 3 h post injection. Scale bar equals 100 µm. **C**-The number of neutrophils recruited to the footpad were scored and expressed as fold change with respect to the number of neutrophils at 1 h post-inflammation. Data are plotted as Means ± SEM with N=3 mice per condition. Two-way ANOVA analysis using Sidak’s multiple comparisons test was used to determine statistical significance. **(D-K)** Neutrophils purified from *WT* or *Alox5*^*-/-*^ or *Blt1*^*-/-*^ mice were labeled with CellTracker Green CMFDA and adoptively transferred to *Alox5*^*-/-*^ **(D-G)** or *Blt1*^*-/-*^ **(H-K)** mice with inflamed footpad. 2P-IVM time-lapse were acquired 1.5 h post *E. coli* BPs injection (Movies S3 and S4). Still images represent maximum intensity projections of Z-stacks in *Alox5*^*-/-*^ **(D)** or *Blt1*^*-/-*^ **(H)** mice. Dashed tangerine lines indicate vessel boundaries (Blue) and magenta arrowheads point to extravasating neutrophils (Green). Scale bar equals 50 µm. Neutrophils were scored for rolling **(E and I)**, arrest **(F and J)** and extravasation **(G and K)** response in *Alox5*^*-/-*^ and *Blt1*^*-/-*^ mice respectively, as described in the Methods section. Data are plotted as % of neutrophils exhibiting a specific behavior and represented as Means ± SEM. Each dot represents data from an individual animal. One-way ANOVA analysis using Dunnett’s multiple comparison test was used to determine statistical significance.

### The LTB_4_-BLT1 axis promotes polarized redistribution of NMIIA and Itgb2 during neutrophil arrest in vivo

The actin-based motor NMIIA is a downstream target of LTB_4_ signaling in neutrophils stimulated with primary chemoattractants (Afonso et al., 2012; Subramanian et al., 2018). To determine how NMIIA activity contributes to neutrophil extravasation, we injected *Alox5*^*-/-*^ mice with *WT* neutrophils pretreated with either Y27632, which inhibits Rho Kinase (ROCK), a key enzyme that controls the activation and assembly of NMIIA filaments (Amano et al., 1996), or (S)-nitro-Blebbistatin (nBleb), which inhibits myosin II contractile activity (Kovács et al., 2004; Lucas-Lopez et al., 2005). We observed that pretreatment with either Y27632 or nBleb, but not the vehicle, significantly reduced both arrest and extravasation in the first 45 min after neutrophil transfer (Fig.2, A-C), consistent with a reported requirement of NMIIA for neutrophil extravasation *in vivo* (Zehrer et al., 2018). However, after 45 minutes, as the inhibitors were cleared, both neutrophil arrest and extravasation resumed similar to vehicle controls, indicating that the effects we observed were not due to drug toxicity (Fig.S1, B-D). These findings led us to address whether the LTB_4_-BLT1 axis regulates NMIIA dynamics during neutrophil extravasation *in vivo*. To this end, neutrophils purified from a knock-in mouse expressing *GFP-NMIIA* (Milberg et al., 2017) were pre-treated with either vehicle or MK886, a covalent non-reversible inhibitor of the 5-lipoxygenase adapter protein (FLAP) (Gillard et al., 1989). First, we determined that the extent of arrest and extravasation of the vehicle-treated *GFP-NMIIA* neutrophils was consistent with that of vehicle-treated *WT* neutrophils (Figs. 2, D-F vs A-C). Second, we found that treatment with MK886 led to reduced arrest and extravasation of *GFP-NMIIA* neutrophils (Fig.2, D-F), consistent with our observations using *Alox5*^*-/-*^ neutrophils. Next, we used ISMic to visualize GFP-NMIIA dynamics in neutrophils that displayed arrest within the vasculature (Movie S5), and score for its cellular distribution during the arrest response. We found that while NMIIA redistributed from the cytoplasm to the cell cortex in the vehicle-treated neutrophils, this response was impaired upon MK886 treatment (Figs. 2G, 2H and S1E; Movie S5). This finding established that LTB_4_ signaling in neutrophils is required for the persistent polarized redistribution of NMIIA *in vivo*. We also attempted to address the impact of the LTB_4_-BLT1 axis on F-actin dynamics during extravasation using neutrophils derived from a mouse expressing the F-actin probe GFP-Lifeact (Riedl et al., 2010). Unfortunately, although *GFP-Lifeact* neutrophils exhibited rolling and arrest, they were defective in extravasation (not shown), suggesting a possible inhibitory effect of this probe. Overall, we conclude that the LTB_4_-BLT1 axis promotes sustained cortical redistribution of NMIIA, which augments neutrophil arrest and extravasation *in vivo*.

**Figure 2.**
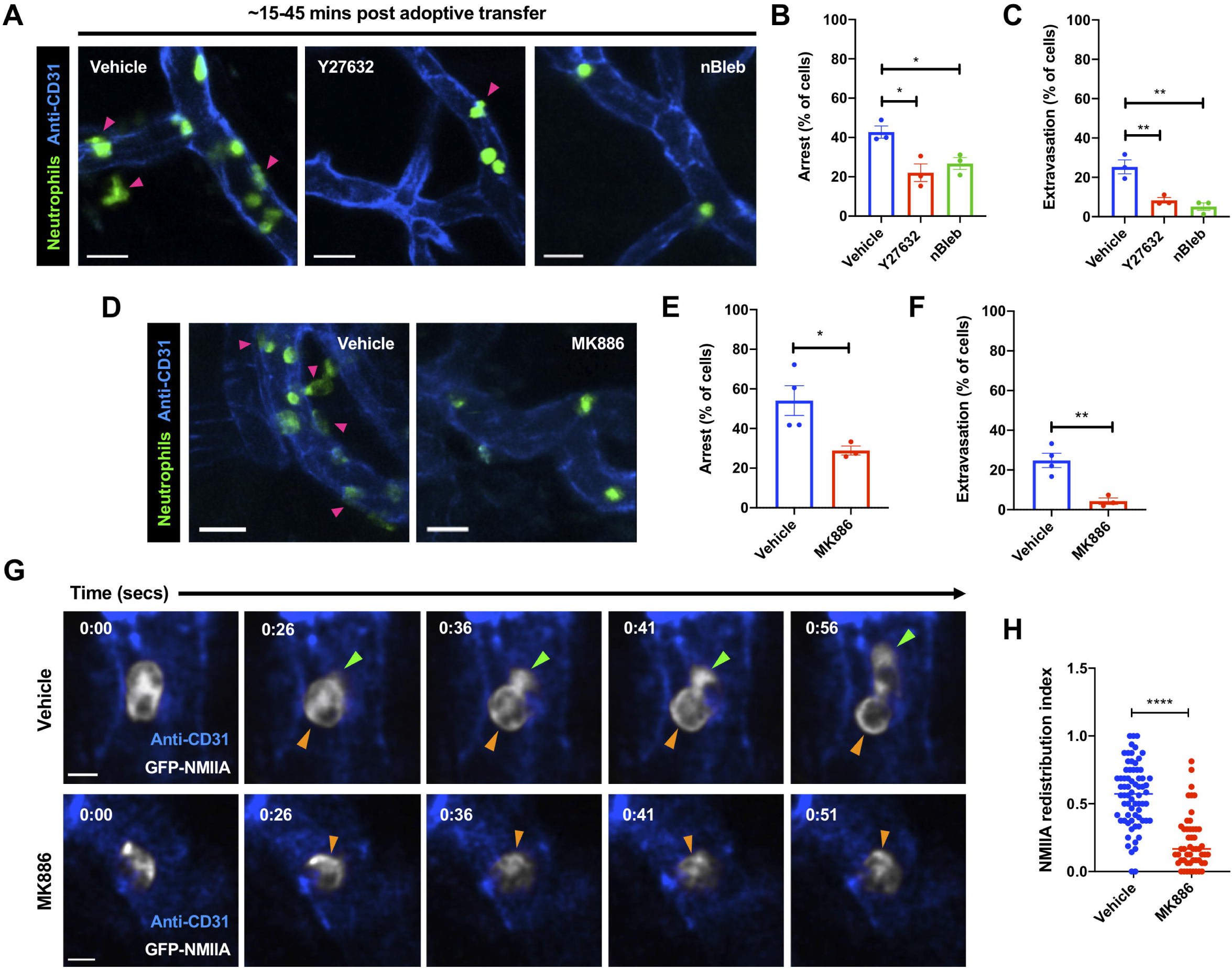
Cortical redistribution of NMIIA during neutrophil arrest requires the LTB_4_-BLT1 signaling axis. **(A-C)** Neutrophils purified from *WT* mice were stained with CellTracker Green CMFDA, treated with 40 µM Y27632 or 10 µM nBleb or the vehicle (DMSO) for ∼30 mins, adoptively transferred into *Alox5*^*-/-*^ mice with inflamed footpad and imaged by 2P-IVM. **A**-Maximum intensity projections of Z-stacks are presented. Magenta arrowheads highlight neutrophils (green) extravasating from the blood vessels (blue). Scale bar equals 20 µm. **B-C** Quantification of the % of neutrophils that exhibited arrest **(B)** or extravasation **(C)** during the first 15-45 mins after neutrophil transfer. Data are plotted as Means ± SEM from N=3 mice for each condition. Unpaired *t* test with Welch’s correction was used to determine statistical significance. **(D-H)** Neutrophils purified from *GFP-NMIIA* mice were treated with either 5 µM MK886 or the vehicle (DMSO), adoptively transferred into *Alox5*^*-/-*^ mice with inflamed footpad and imaged by ISMic. **D**-Maximum intensity projections of Z-stacks are presented. Magenta arrowheads highlight extravasating *GFP-NMIIA* neutrophils (green) from the blood vessels (blue). Scale bar equals 20 µm. **(E-F)** Quantification of the % of neutrophils that exhibited arrest **(E)** or extravasation **(F)** response. Data are plotted as Means ± SEM, N=4 mice for vehicle and N=3 mice for the MK886 treatment. Unpaired *t* test with Welch’s correction was used to determine statistical significance. **G**-Still images represent an individual optical slice from a Z-stack acquired in time-lapse (see Movie S5). GFP-NMIIA is in white and blood vessels in blue. Green arrowhead points to the protrusive front of the extravasating neutrophil. Orange arrowheads indicates a region of cortical NMIIA enrichment. Time is represented in mins:secs. Scale bar equals 5 µm. **H**– The cortical redistribution of NMIIA was determined as described in the Methods section and in Fig. S1E and represented as redistribution index. Data points represent 70 cells from N=4 mice for the vehicle and 51 cells from N=3 mice for MK886 treatment, with each dot representing value of a single cell. Unpaired *t* test with Welch’s correction was used to determine statistical significance.

Although NMIIA likely provides the contractile force required for neutrophils to squeeze through extravasation sites, its roles during arrest is less clear. Therefore, we further investigated this aspect *in vivo*. Leukocyte arrest in the inflamed vessels, in most tissues, requires the engagement of Itgb2 on the plasma membrane (PM) with inter cellular adhesion molecules (ICAMs) presented on the inflamed endothelium (Liew and Kubes, 2019). Consistent with this notion we found that *Itgb2*^*-/-*^ neutrophils, when compared to *WT* neutrophils, displayed increased rolling as well as severe defects in arrest and extravasation when introduced in *Alox5*^*-/-*^ mice with inflamed footpad (Fig.3, A and B), similar to our observations with *Alox5*^*-/-*^ neutrophils. Also, LTB_4_-stimulated neutrophils utilize Itgb2 on the PM to engage with ICAM1-coated surface *in vitro* (Colom et al., 2015). We therefore hypothesized that the LTB_4_-BLT1-NMIIA pathway regulates the dynamics and localization of Itgb2 at the PM to promote firm neutrophil arrest. To directly visualize Itgb2 dynamics, we incubated *WT* or *Alox5*^*-/-*^ neutrophils with a fluorescently-conjugated antibody to Itgb2 (M18/2 clone) and introduced them into *Alox5*^*-/-*^ mice (Fig.3C). The antibody bound specifically to Itgb2 and did not qualitatively impact the extravasation response of labeled *WT* neutrophils *in vivo* (Fig.3, C and D; Movie S6). In *WT* neutrophils that rolled along the blood vessel, Itgb2 localized in cytoplasmic clusters (Fig.3D, left panel and Movie S6). As the *WT* neutrophil underwent arrest, Itgb2 redistributed to areas of the cell periphery that were in direct contact with the endothelium (Fig.3D, middle panel and Movie S6). Later, as the *WT* neutrophil protruded and extravasated out of the vessel, Itgb2 gradually redistributed to the back of the migrating cell (Fig.3D, right panel and Movie S6), consistent with the previously observed high-affinity and clustering patterns of Itgb2 isoforms during neutrophil extravasation *in vivo* (Hyun et al., 2019). Conversely, Itgb2 remained dispersed in the cytoplasm and failed to redistribute to the cell periphery in *Alox5*^*-/-*^ neutrophils that transiently underwent arrest response (Fig.3, E and F; Movie S6). A similar defect in Itgb2 redistribution was also observed in *WT* neutrophils that were pre-treated with either Y27632 or nBleb (Fig.3F), indicating a requirement for NMIIA in this process. These findings suggest that NMIIA activation downstream of LTB_4_ signaling facilitates the dynamic redistribution and sustained localization of Itgb2 on the neutrophil PM to promote arrest response in the inflamed endothelium.

**Figure 3.**
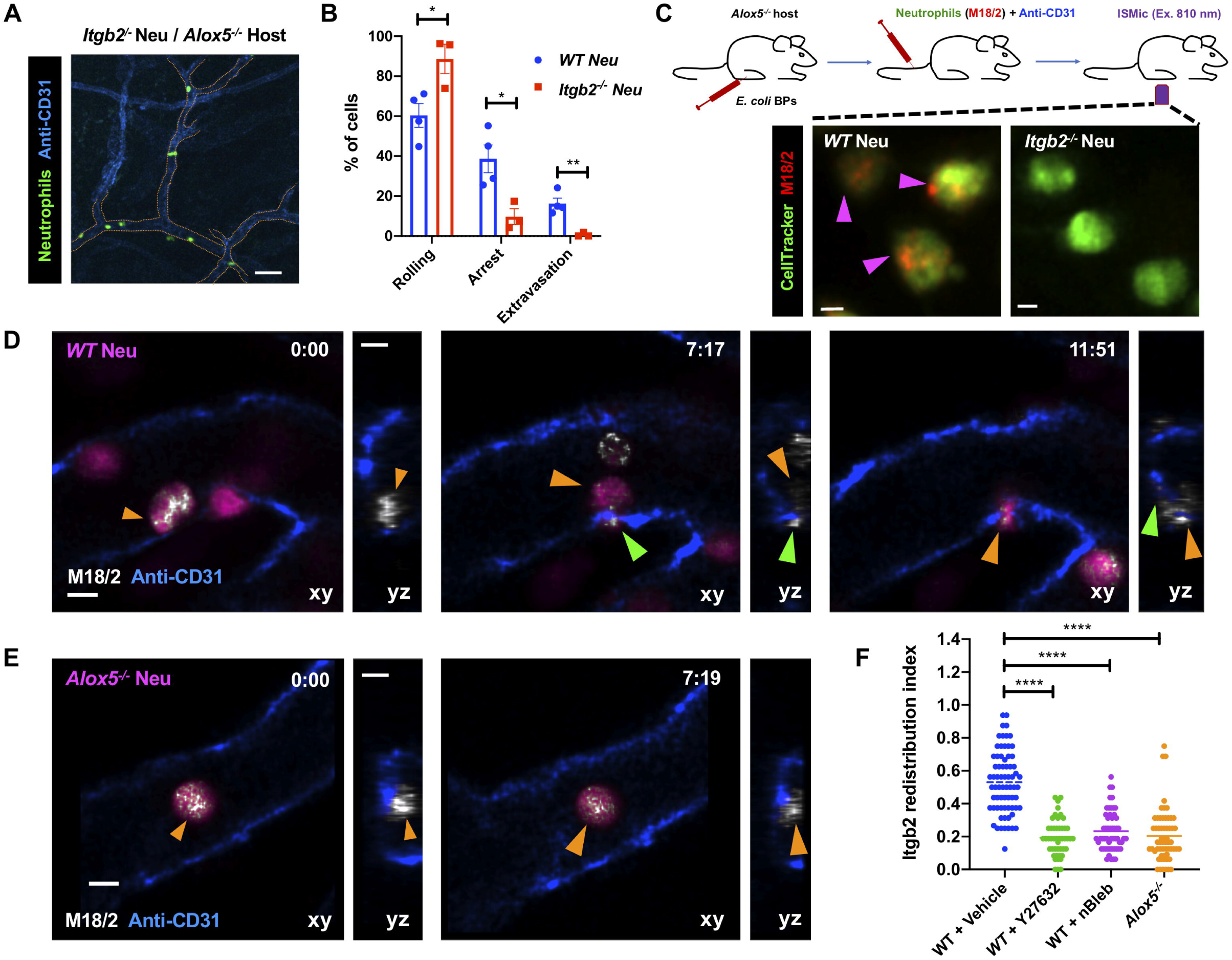
LTB_4_-BLT1 axis regulates Itgb2 dynamics during neutrophil extravasation *in vivo*. **(A-B)** Neutrophils purified from *Itgb2*^*-/-*^ mice were labeled with CellTracker Green CMFDA, adoptively transferred into *Alox5*^*-/-*^ mice with inflamed footpad and imaged by 2P-IVM. **A**-Maximum intensity projection of a Z-stack. Dashed tangerine lines indicate vessel boundaries. Scale bar equals 50 µm. **B** – Quantitative analysis of the % of neutrophils that exhibit rolling, arrest and extravasation response in *Itgb2*^*-/-*^ neutrophils is compared to its *WT* counterpart. Values for *WT* neutrophils are the same as in Fig. 1E-G. Data are plotted as Means ± SEM and each dot represents result from an individual animal. Unpaired *t* test with Welch’s correction was used to determine statistical significance. **(C)** Neutrophils purified form either *WT* or *Itgb2*^*-/-*^ mice were labeled with CellTracker Green CMFDA and a fluorescently-conjugated antibody against mouse Itgb2 (M18/2 clone, red), adoptively transferred into *Alox5*^*-/-*^ mice with inflamed footpad and imaged by ISMic (top panel). Maximum intensity projections of Z-stacks are presented (bottom panels). Magenta arrowheads indicate neutrophils positive for Itgb2 staining. The results represent N=3 mice per condition. Scale bar equals 3 µm. **(D-F)** Neutrophils purified form either *WT* or *Alox5*^*-/-*^ mice were labeled with CellTracker Green CMPTX and fluorescently-conjugated M18/2 antibody (red), adoptively transferred into *Alox5*^*-/-*^ mice with inflamed footpad and imaged by ISMic. Additionally, *WT* neutrophils were pre-treated with either vehicle (DMSO; **D** and **F**) or 40 µM Y27632 (**F**) or 10 µM nBleb (**F**) and transferred into the recipient. Still images represent maximum intensity projections in xy (large panels) and zy (side panels) dimensions, derived from time-lapse sequences in Movie S6 (mins:secs) for *WT* neutrophils (**D**) and *Alox5*^*-/-*^ neutrophils (**E**). CellTracker is colored in magenta, blood vessels in blue and anti-Itgb2 in white. Orange and green arrowheads indicate the back and protrusive front of arrested and extravasating neutrophil. Scale bars equal 5 µm. **F**– The Itgb2 redistribution index was measured as described in the Methods section and presented as individual points for individual cells of *WT* (DMSO treated; 70 cells), *WT* + Y27632 (47 cells), *WT* + nBleb (54 cells) and *Alox5*^*-/-*^ (59 cells), from N=3 mice per condition. One-way ANOVA analysis using Dunnett’s multiple comparison test was used to determine statistical significance.

### The LTB_4_-BLT1 axis promotes adhesion, actomyosin polarization and Itgb2 trafficking in primary neutrophils in vitro

LTB_4_ has been shown to act as a signal-relay molecule to promote chemotaxis of primary human neutrophils (PMNs) in response to primary chemoattractant (Afonso et al., 2012). Since we found that the LTB_4_-BLT1 axis is required for neutrophil arrest in mice, we asked if such a mechanism operates during PMN adhesion in response to primary chemoattractant stimulation. To this end, we analyzed stimulation-induced adhesion of PMNs on a fibrinogen-coated surface. PMNs pre-treated with vehicle showed a progressive increase in their ability to adhere upon stimulation with the bacterial peptide analogue fNLFNYK (Fig.4A). However, pre-treatment with MK886 or LY223982, a BLT1 antagonist (Jackson et al., 1992), significantly reduced PMN adhesion at later time points (Fig.4A). Importantly, the requirement of the LTB_4_-BLT1 axis for sustained PMN adhesion was specific to end-target (fNLFNYK and C5a), but not intermediate (CXCL8) chemoattractants (Fig. S2A), consistent with the notion that the LTB_4_-BLT1 axis functions downstream of end-target or primary chemoattractants (Afonso et al., 2012; Sadik et al., 2012; Subramanian et al., 2018). Similarly, mouse neutrophils from *Blt1*^*-/-*^ mice adhered significantly less when compared to their *WT* counterparts upon stimulation with the bacterial peptide analogue, WKYMVm (Fig.S2B). Moreover, neutrophils from *GFP-NMIIA* mice adhered to a similar extent as *WT* controls upon WKYMVm stimulation (Fig.S2C). However, treatment with MK886 or LY223982, as compared to the vehicle control, significantly reduced the extent of adhesion of *GFP-NMIIA* neutrophils (Fig.S2D), as we observed in PMNs. Importantly, MK886 or LY223982 treatment reduced the extent of cortical GFP-NMIIA and polarized F-actin distribution in *GFP-NMIIA* neutrophils in response to WKYMVm stimulation (Fig.S2, E and F). Similarly, a defect in cortical actomyosin distribution was also observed in PMNs upon treatment with MK886 or LY223982 (Fig.4, B-D). Therefore, the LTB_4_-BLT1 axis relays signals to promote sustained actomyosin polarity and adhesion of primary neutrophils downstream of primary chemoattractant stimulation.

**Figure 4.**
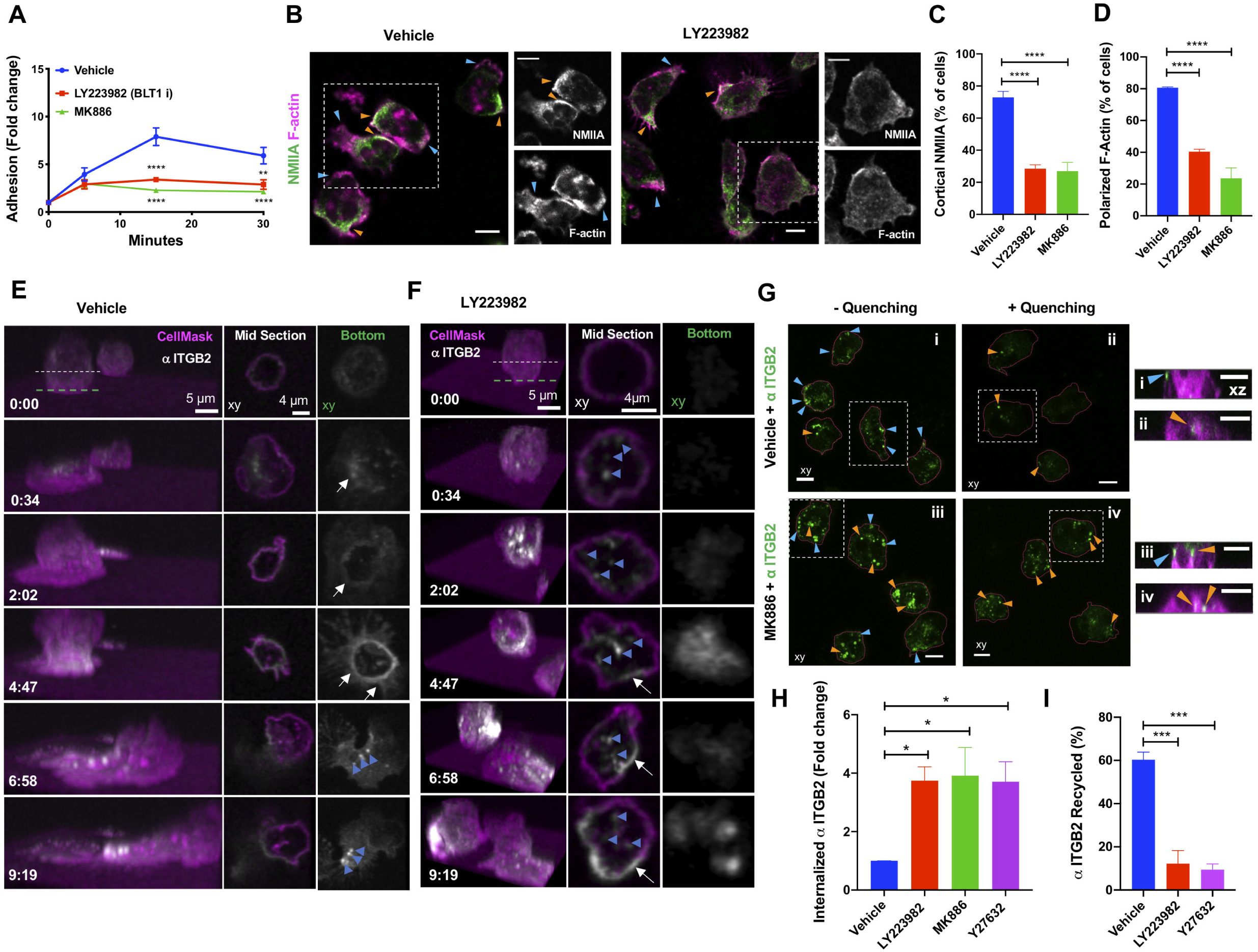
PMNs require LTB_4_-BLT1 axis for sustained polarization, ITGB2 trafficking and adhesion response *in vitro*. **(A)** PMNs were pre-treated for 20 mins with vehicle or 2 µM MK886 or 20 µM LY223982 and stimulated with 25 nM fNLFNYK on a fibrinogen-coated surface. Cells remained either unstimulated (30 mins) or were stimulated for 30, 15 and 5 mins independently prior to fixation. Adhesion was calculated as described in the Methods section and presented as fold change of adhered neutrophils in comparison to unstimulated controls. Data are plotted as Means ± SEM from N=3 independent experiments. Two-way ANOVA analysis using Dunnette’s multiple comparison test was used to determine statistical significance. **(B-D)** PMNs were pre-treated with the vehicle or 2 µM MK886 or 20 µM LY223982 for 20 mins, stimulated with 25 nM fNLFNYK for 15 mins, fixed and stained with rhodamine phalloidin (magenta) and anti-NMIIA (green). **B**-Representative confocal images for vehicle and LY223982 treated PMNs. Blue and orange arrowheads indicate the front and back of PMNs with polarized actomyosin distribution. Dashed white box indicates the region zoomed and presented on the right side for each condition, with individual channels in gray scale. Scale bars equal 5 µm. **C-D** Quantification of the % neutrophils exhibiting cortical NMIIA **(C)** and polarized F-actin **(D)** in response to above mentioned treatments. Data are plotted as Means ± SEM from N=3 independent experiments. One-way ANOVA analysis using Dunnette’s multiple comparison test was used to determine statistical significance. **(E-F)** PMNs were pre-treated with either the vehicle **(E)** or 20 µM LY223982 **(F)** for 20 mins. They were incubated with an Alexa488-conjugated antibody against human ITGB2 (αITGB2, CTB104 clone; white) in combination with CellMask Deep Red (magenta) for ∼ 1 min before stimulation with 100 nM fNLFNYK for 10 mins on a fibrinogen-coated surface and acquired in 3D using time-lapse confocal microscopy. See Movie S7. Left panel represents the 3D view of the cell as a function of time (mins:secs). Middle and right panels represent the middle and bottom slices, as indicated by the dashed lines in the 3D view respectively. White arrows indicate the formation of ITGB2 ring-like structure. Blue arrowheads indicate ITGB2 vesicles. Images are representative of 3 independent experiments (see Figs. S2G-I). **(G)** PMNs were labeled with CellMask Deep Red to visualize the PM (magenta) and with αITGB2 (green), pre-treated with either the vehicle or 2 µM MK886 for 20 mins and stimulated with 25 nM fNLFNYK on a fibrinogen-coated surface. Neutrophils were fixed, and either left untreated (-quenching; i and iii) or treated with an anti-Alexa Fluor 488 antibody (+quenching; ii and iv). Images represent the maximum intensity projections in xy (large panels on the left) dimension for each condition. The cells in dashed white boxes are represented in the xz dimension in small panels on the right. Blue and orange arrowheads indicate surface and internalized anti-ITGB2 signals respectively. Scale bars equal 5 µm. Images are representative of N=3 independent experiments. **(H-I)** Internalization and recycling of ITGB2. **H**-PMNs were treated with the vehicle or 2 µM MK886 or 20 µM LY223982 or 20 µM Y27632, for 20 min and stimulated with 25 nM fNLFNYK for 10 min on a fibrinogen-coated surface in the presence of αITGB2. Samples were processed and imaged as described in the Methods section to determine the % of the internalized ITGB2. Data are plotted as Means ± SEM from N=4 independent experiments. **I**-PMNs were treated with the vehicle or 2 µM MK886 or 20 µM LY223982 or 20 µM Y27632, for 20 min and stimulated with 25 nM fNLFNYK for 15 min on a fibrinogen-coated surface in the presence of αITGB2. Samples were washed and either fixed or stimulated for additional 45 mins in the presence of fNLFNYK and inhibitors, before fixation. Samples were processed and imaged as described in the Methods section to determine the extent of recycling of internalized Itgb2. Data are plotted as Means ± SEM from N=3 independent experiments. One-way ANOVA analysis using Dunnette’s multiple comparison test was used to determine statistical significance.

Although we employed ISMic in the past to track endocytic events in other organ systems (Masedunskas and Weigert, 2008; Shitara et al., 2019), due to the depth of imaging in the mouse footpad, we could not image the trafficking of the Itgb2-containing intracellular structures with sufficient temporal and spatial resolution *in vivo*. Therefore, we used time-lapse confocal imaging to study the dynamics of human Itgb2 (ITGB2) in PMNs. To this end, we used an Alexa 488-conjugated antibody to label ITGB2 (CTB104 clone, hereafter called fluorescent αITGB2) in PMNs. We found that, as PMNs began spreading upon fNLFNYK stimulation, ITGB2 quickly redistributed along the PM to form a ring-like structure at the interface of PMN contact with the fibrinogen-coated surface (Fig.4E white arrows and Movie S7A), reminiscent of our observations using ISMic (Fig.3) and with previous *in vitro* findings (Rochon et al., 2000; Shaw et al., 2004). With time, the ring-like structure disassembled, and ITGB2-containing vesicles internalized from the back of polarized and migrating PMNs (Fig.4E, blue arrowheads and Movie S7A). However, upon blocking LTB_4_ signaling using LY223982, we observed an increase of ITGB2-containing vesicles, rather than the formation of the ring-like structure (Fig.4F, blue arrowheads and Movie S7B). Consistently, a significant increase in the extent of internalized ITGB2-containing vesicles upon LY223982 treatment was also observed (Fig.S2, G-I). This finding prompted us to further investigate the regulation of ITGB2 trafficking by the LTB_4_-BLT1 axis in PMNs. In polarized vehicle-treated PMNs, we found that clusters of ITGB2 largely localized at the PM around the adhesion site (Fig.4G, i), as revealed by the quenching of fluorescence of the αITGB2 antibody following incubation with a cell-impermeant antibody directed against the fluorophore (Fig.4G, ii). On the other hand, in MK886-treated PMNs most of the αITGB2 fluorescence showed a punctate intracellular distribution, which persisted even after treatment with the quenching antibody (Fig.4G, iii and iv). Using inhibitors to specific endocytic pathways, we confirmed that the internalization of αITGB2 indeed occurred via a clathrin-independent and dynamin-dependent endocytic pathway (Fig.S2, J and K), as previously described (Fabbri et al., 2005). Moreover, treatment with inhibitors of either the LTB_4_-BLT1 axis or ROCK, increased the pool of internalized ITGB2 with respect to control (Fig.4H). Finally, the extent of increase in the internalized ITGB2 corresponded to a block in the recycling of ITGB2 back to the PM, since only ∼15% of the internalized ITGB2 was cleared upon inhibition of the LTB_4_-BLT1-myosin pathway, compared to 60% in the control (Fig.4I). Together, these findings in PMNs unravel a role for the LTB_4_-BLT1-myosin axis in promoting recycling of ITGB2 from an intracellular pool as well as its retention on the PM (Fig.S2L), which is consistent with our ISMic analysis in live mice.

Overall, we found that NMIIA regulates integrin-based adhesion stability during neutrophil arrest, possibly via different non-exclusive mechanisms, as previously reported (Vicente-Manzanares et al., 2009). We document that Itgb2 redistributes from a dispersed intracellular pool to a defined area of the neutrophil surface that is juxtaposed to the inflamed endothelium (adhesion ring), consistent with previous *in vivo* and *in vitro* studies (Hyun et al., 2019; Jones et al., 1988; Shaw et al., 2004). At this stage, we cannot resolve whether Itgb2 clusters observed at the adhesion ring-like structures formed through lateral diffusion or rapid internalization and recycling back to the PM. We favor the latter explanation as inhibition of the LTB_4_-BLT1 axis resulted in: (i) a lack of the Itgb2 adhesion ring-like structure; (ii) an increased Itgb2 internalization; and (iii) an impaired Itgb2 recycling to the PM. Notably, these defects were phenocopied upon inhibition of NMIIA activation, thus strongly linking NMIIA activity with Itgb2 trafficking in neutrophils. Whether NMIIA activation directly promotes Itgb2 recycling or negatively regulates its internalization has yet to be determined. However, the regulation of Itgb2 by NMIIA *in vivo* is likely indirect, since they are localized to the front and rear of arrested neutrophils respectively. As NMIIA activity is proposed to regulate lipid microdomains on the neutrophil PM (Hind et al., 2016), it is conceivable that NMIIA, at the rear of the cells, creates domains that limit Itgb2 internalization and force the rapid recycling of internalized Itgb2 to the front of cells, thus confining its localization to adhesion sites on the PM. However, in the absence of the LTB_4_-BLT1 axis, NMIIA controlled microdomains might be impaired, resulting in enhanced internalization and reduced oriented Itgb2 recycling, likely diverting its trafficking to a slow recycling pathway. The net result would be a reduction in Itgb2 levels on the PM, leading to an impairment in arrest, and consequently in neutrophil extravasation.

### The autocrine/paracrine action of LTB_4_ requires EV release from neutrophils

As LTB_4_ produced by neutrophils drives their arrest and extravasation *in vivo*, we investigated whether this process occurs in an autocrine/paracrine fashion. If this is the case, we reasoned that defects in arrest and extravasation observed in *Alox5*^*-/-*^ neutrophils should be rescued upon co-injection with *WT* neutrophils, which are capable of producing LTB_4_ (Fig.S3, A and B). Indeed, *Alox5*^*-/-*^ neutrophils displayed robust rolling, arrest, and extravasation in the presence of vehicle-treated *WT* neutrophils (Fig.5, A-i and B; Movie S8), but not with MK886-treated *WT* neutrophils (Fig.5, A-ii and B; Movie S8). We next investigated whether LTB_4_ release and signal-relay occurred via EVs, as reported during neutrophil chemotaxis *in vitro* (Majumdar et al., 2016). Using ISMic we observed large Itgb2-containing neutrophil-derived EVs, which rolled along and occasionally tethered to the inflamed endothelial surface facing the vascular lumen in the vicinity of extravasating neutrophils (Fig.5, C and D; Movie S9). Due to the limits in resolution of light microscopy, we could not determine whether these large vesicles were shed as individual membranous compartments or clusters of smaller vesicles. Therefore, we examined whether their biogenesis and release shared any common machinery with the generation of EVs, which in neutrophils are regulated by both the neutral sphingomyelinase (N-SMase) and the small GTPase Rab27a (Majumdar et al., 2016). We found that the number of neutrophil-derived EVs in the vasculature was dramatically reduced in *WT* neutrophils pre-treated with either the N-SMase inhibitor GW4869 (Luberto et al., 2002) or the Rab27a inhibitor Nexinhib20 (Johnson et al., 2016) (Fig. 5E). On the other hand, blocking NMIIA contractility (nBleb), lack of LTB_4_ production (*Alox5*^*-/-*^) or Itgb2 (*Itgb2*^*-/-*^), did not impact the number of EVs observed in the vasculature (Fig.5, E and F).

**Figure 5.**
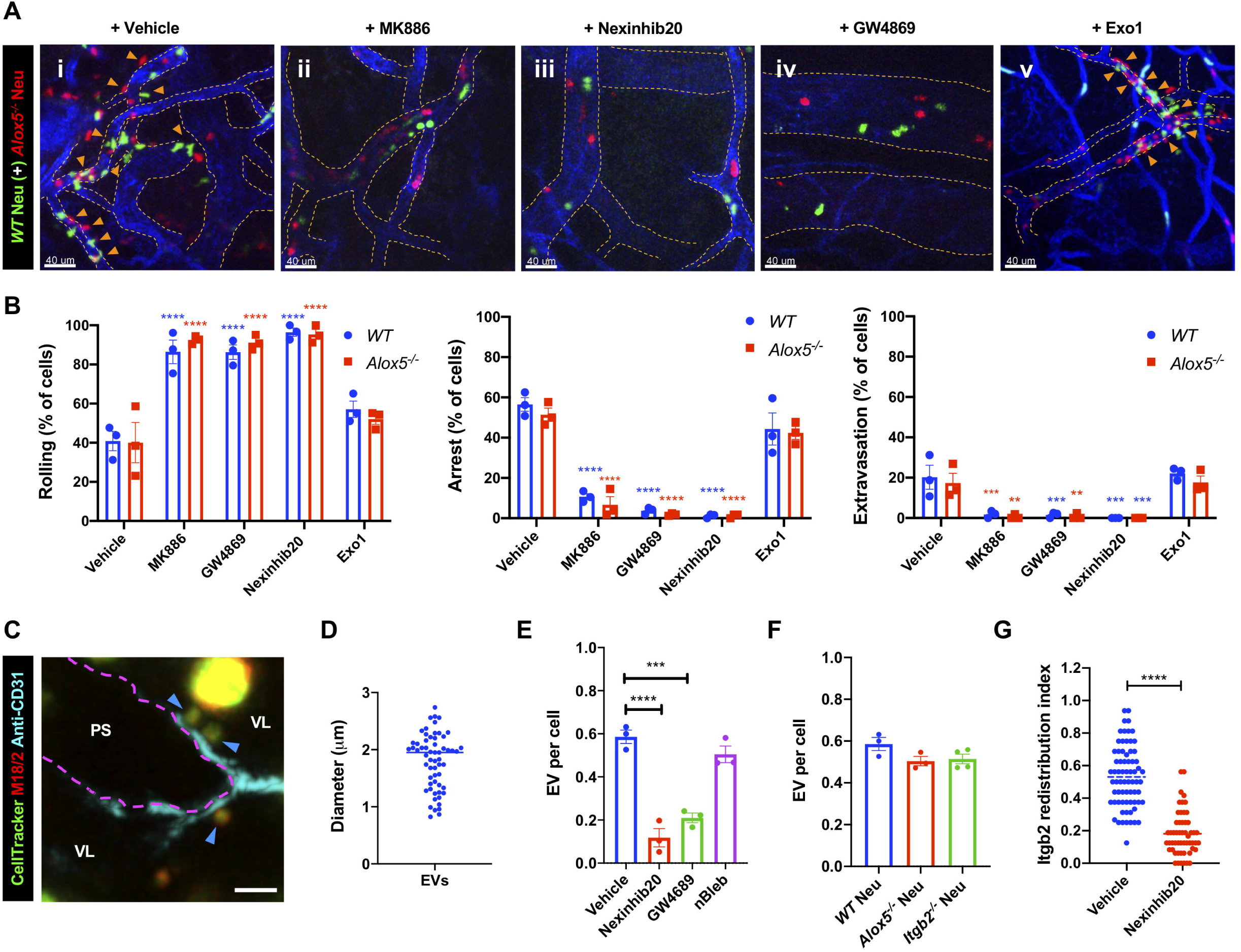
EVs mediate the autocrine/paracrine action of LTB_4_ during neutrophil arrest and extravasation response. **(A-B)** *WT* neutrophils labeled with CellTracker Green CMFDA (green), were pre-treated with vehicle (DMSO; i) or 5 µM MK886 (ii), 2 µM Nexinhib20 (iii), 20 µM GW4689 (iv) or 40 µM Exo1 (v) for 30 mins, and co-injected into *Alox5*^*-/-*^ mice with inflamed footpad, and *Alox5*^*-/-*^ neutrophils labeled with CellTracker Red CMPTX (red). **A**-Maximum projections of Z-stacks acquired by 2P-IVM. See Movie S8. Blood vessels are in blue and dashed orange lines indicate vessel boundaries. Orange arrowheads point to extravasated neutrophils. Scale bar equals 40 µm. **B-** Analysis of % of neutrophils displaying rolling, arrest and extravasation in the above-mentioned conditions. Data are plotted as Means ± SEM from N=3 mice per group. Two-way ANOVA analysis using Dunnett’s multiple comparison test was used to determine statistical significance. **(C-D)** *WT* neutrophils were labeled with CellTracker Green CMFDA (green) and fluorescently conjugated M18/2 antibody (Red), before their adoptive transfer into *Alox5*^*-/-*^ mice with inflamed footpad and imaged by ISMic. **C-** A representative maximum projection of a Z-stack with blood vessel in cyan (boundary marked with magenta dotted line), blue arrowheads indicate the EVs that rolled and tethered to the inflamed endothelial surface facing the vascular lumen (VL), and ‘PS’ indicates the perivascular space. Scale bar equals 5 µm. See Movie S9. **D-** Quantification of the size of EVs detected using ISMic analysis. Each dot represents single EV and a total of 60 EVs were analyzed from N=3 independent experiments. **(E-F)** Neutrophils purified form *WT* **(E and F)** or *Alox5*^*-/-*^ **(F)** or *Itgb2*^*-/-*^ **(F)** mice were labeled with CellTracker Green CMPTX and fluorescently-conjugated M18/2 antibody (red), adoptively transferred into *Alox5*^*-/-*^ mice with inflamed footpad and imaged by ISMic. Additionally, *WT* neutrophils were pre-treated with either vehicle (DMSO) or 2 µM Nexinhib20 or 20 µM GW4689 or 10 µM nBleb for 30 mins, before their adoptive transfer into the recipient. The number of EVs detected was normalized to the number of detected neutrophils for each condition, as described in the Methods section. Each dot represents the result from an individual animal experiment. Data are plotted as Means ± SEM. One-way ANOVA analysis using Dunnette’s multiple comparison test was used to determine statistical significance and no significant difference was observed in panel **F**. **(G)** Neutrophils purified form *WT* mice pre-treated with either vehicle (DMSO) or 2 µM Nexinhib20 for 30 mins, labeled with CellTracker Green CMPTX and fluorescently-conjugated M18/2 antibody (red), adoptively transferred into infected *Alox5*^*-/-*^ mice with inflamed footpad and imaged by ISMic. The peripheral Itgb2 redistribution index was determined as described in the Methods section. Each dot represents the value for an individual cell, with values for *WT* vehicle control the same as in Fig. 3F and n=54 cells for Nexinhib20, from N=3 mice for each group. Unpaired *t* test with Welch’s correction was used to determine statistical significance.

Based on these findings, we speculated that EVs could mediate the autocrine/paracrine actions of LTB_4_ (Fig.S3C). This hypothesis was confirmed by the fact that *WT* neutrophils pre-treated with either GW4869 (Feng et al., 2003) or Nexinhib20, but not the control drug Exo1, failed to rescue the defect in arrest and extravasation of *Alox5*^*-/-*^ neutrophils (Fig.5, A-i, iii-v and B; Movie S8). Consistent with these results, we found that the inhibitors of EV release reduced actomyosin polarization, enhanced ITGB2 internalization and impaired adhesion in PMNs stimulated with fNLFNYK *in vitro* (Fig.S3, D-G). Moreover, treatment with Nexinhib20 impaired the redistribution of Itgb2 in *WT* neutrophils *in vivo* (Fig.5G). Together, these findings suggest that neutrophil arrest and extravasation *in vivo* is mediated via EVs, which facilitate LTB_4_ relay in an autocrine/paracrine manner (Fig.S4). This is, to our knowledge, the first evidence of an autocrine/paracrine LTB_4_ signaling mechanism in vascular neutrophils that requires EVs, a mechanism in line with the emerging paradigm on the ability of EVs to relay chemotactic signals during amoeboid cell migration (Kriebel et al., 2018; Majumdar et al., 2016). Given the limitations of light microscopy, we document large neutrophil-derived EVs (size ∼1-2 µm) in the vasculature, similar to the EVs documented from neutrophils and tumor cells *in vivo* (Hoshino et al., 2015; Lai et al., 2015; Lim et al., 2015). The EVs we observed are similar to the microvesicles observed from the trailing-edge of trans-migrating neutrophils that get deposited in the perivascular area (Hyun et al., 2012). However, while the microparticles in trans-migrating neutrophils have been suggested to require Itgb2 (Hyun et al., 2012), the generation of EVs we observed did not require Itgb2 in neutrophils. Interestingly, the release of neutrophil-derived EVs in our model system requires N-SMase and Rab27 activity, which have previously been described to regulate the release of small EVs (Catalano and O’Driscoll, 2019; Majumdar et al., 2016; Raposo and Stoorvogel, 2013). However, whether such EVs harbor the LTB_4_-synthesizing machinery and to what extent they utilize Itgb2 to tether to the inflamed endothelium remains to be ascertained.

In this study, we used two-photon intravital microscopy (2P-IVM) to investigate the specific mechanism by which the LTB_4_-BLT1 axis signals during intravascular neutrophil response in the early phase of bacterial bioparticle-induced inflammation. Using pharmacological and genetic tools, we found that neutrophils undergo firm arrest and extravasation by producing and sensing LTB_4_ in an autocrine/paracrine manner utilizing an EV-based mechanism. Mechanistically, LTB_4_ signaling is required for sustained actomyosin-based polarity, which ensures oriented vesicle trafficking and Itgb2 expression at the PM to promote persistent and productive interaction with the inflamed endothelium. While neutrophils appear to be the major source of LTB_4_ in our model system, it is conceivable that the LTB_4_ produced from resident non-neutrophil cell types can contribute to the paracrine signaling under other physiological conditions.

An important aspect we haven’t addressed in this study, and is currently under investigation, is that of primary signals that initiate the LTB_4_ cascade in neutrophils. Several mechanisms applicable to our model system could be occurring, such as exposure of tissue to: (i) primary chemoattractants, which are sufficient to induce neutrophil extravasation from the exposed vessels (Hyun et al., 2019; Miyabe et al., 2017) through atypical receptors (Girbl et al., 2018; Miyabe et al., 2019); (ii) bacteria, which stimulate perivascular macrophages to provide chemoattractant cues to direct neutrophil extravasation in dermal venules (Abtin et al., 2014); and (iii) tumor necrosis factor α (TNFα) (Camussi et al., 1989) or bacterial lipopolysaccharide (Doerfler et al., 1989), which could prime neutrophils to generate LTB_4_, possibly via ATP-gated calcium channels (Poplimont et al., 2020; Schumann et al., 1993). Once stimulated, autocrine/paracrine LTB_4_-BLT1 signaling in neutrophils could likely function in conjunction with CXCL1 present on the inflamed endothelial surface to promote firm neutrophil arrest, and potentiate neutrophil elastase secretion, which would help unseal endothelial junctions to facilitate CXCL2-guided neutrophil trans-migration (Colom et al., 2015; Girbl et al., 2018). We envision a scenario where neutrophils, upon encountering primary signals, release EVs into the vasculature in proximity to the inflamed site. Given that EVs from stimulated neutrophils can utilize Itgb2 to anchor to the extracellular matrix and secrete neutrophil elastase to cleave the matrix (Genschmer et al., 2019), it is tantalizing to speculate that the EVs in the vasculature tether to the inflamed endothelium in an Itgb2-dependent fashion and locally release both LTB_4,_ and neutrophil elastase. LTB_4_ and neutrophil elastase would in turn relay chemotactic signals to promote neutrophil arrest and to degrade endothelial junctions, respectively. Furthermore, as the inflammation proceeds, EVs could persistently engage with the inflamed vessels to serve as ‘beacons’ to attract more neutrophils and other immune cells from the vasculature to follow the neutrophil path to inflamed sites (Lim et al., 2015; Oyoshi et al., 2012). Overall, our findings bear important clinical relevance, as LTB_4_ production and EV generation in neutrophils are intimately associated with the progression of diseases including tumor metastasis (Wculek and Malanchi, 2015) and chronic inflammatory lung disorder (Genschmer et al., 2019), respectively.

## MATERIALS AND METHODS

### Mice

The following strains: *C57BL/6J (WT), Alox5*^*-/-*^ (B6.129S2-*Alox5*^*tm1Fun*^*/J*), *Blt1*^*-/-*^ (B6.129S4-*Ltb4r1*^*tm1Adl*^*/J*) and *Itgb2*^*-/-*^ (B6.129S7-*Itgb2*^*tm1Bay*^*/J*) were obtained from JAX (The Jackson Laboratory, Bar Harbor). *LyzM-GFP, GFP-NMIIA* and *GFP-Lifeact* mice were obtained from Drs. Dorian McGavern (NINDS, NIH), Robert Adelstein (NHLBI, NIH) and Roland Wedlich-S_Ö_ldner (Riedl et al., 2010), respectively. The study was approved and conducted in accordance with the animal protocols approved by the Institutional Animal Care and Use Committee, protocols – LCMB-031 and LCMB-035 (National Cancer Institute). Mice, both males and females, were used for the experiments at an age between 2-6 months.

### Reagents

Antibodies against mouse Itgb2 (clone M18/2; Alexa Fluor 594 conjugated), NMIIA heavy chain (clone Poly19098) and custom Rhodamine-conjugated antibody to CD31 (clone 390) were from BioLegend, Inc. Antibodies against CD31 (clone 390; eFluor450 conjugated) and for quenching Alexa Fluor 488 (RRID – AB_2532697) were from Invitrogen. Antibodies against CD16/32 (Fc block; clone 2.4G2) and ITGB2 (clone CTB104; Alexa Fluor 488 conjugated) were from BD Pharmingen and Santa Cruz Biotechnology Inc., respectively. *E. coli* BPs (K12-strain; Texas red conjugated), CellTracker Green CMFDA, CellTracker Red CMPTX, CellMask Deep Red for PM and Rhodamine Phalloidin dyes were from Molecular Probes. PitStop2, MDC, fibrinogen (human plasma derived) and WKYMVm were from Sigma Aldrich. Recombinant fNLFNYK, human C5a and human CXCL8 were from Santa Cruz Biotechnology Inc., R & D Systems and Peprotech Inc., respectively. LTB_4_, LY223982, Y27632, nBleb and GW4869 were from Cayman Chemical. nBleb used in this study is a photostable version of Blebbistatin (Lucas-Lopez et al., 2005). MK886, Nexinhib20, Exo1, Dynasore and MitMAB were from Tocris Bioscience.

### Primary neutrophil isolation from mice and human samples

Mouse neutrophils were obtained from the bone marrows of front and hind limbs. Briefly, cells were flushed out using RPMI without phenol red medium (Invitrogen) supplemented with 1 mM HEPES (Gibco) and antibiotics (Gibco) (hereafter called RPMI medium) and filtered via a 40 µm cell strainer (Corning Inc.). Subsequently, red blood cells were lysed using ACK buffer (Gibco) for 30 secs, layered onto a discontinuous gradient of Histopaque 1077 (Sigma Aldrich) and Histopaque 1119 (Sigma Aldrich), in a 1:1:1 ratio as previously reported (Swamydas et al., 2015). Neutrophils were isolated from the 1077/1119 Histopaques interface, rinsed and resuspended in RPMI medium. Human neutrophils (PMNs) were obtained from heparinized whole blood of healthy human donors by venipuncture, as part of the NIH Blood Bank research program and purified from 1077 Histopaque-RPMI medium interface post differential centrifugation as reported previously (Subramanian et al., 2018). Red blood cells were then lysed by multiple rounds of exposure to hypotonic solutions (0.2% NaCl solution; 30 secs) followed by the addition of a neutralizing solution (1.6% NaCl solution), and then centrifugation. Isolated primary neutrophils were resuspended in RPMI medium and incubated on a rotator at room temperature (hereafter called RT) for further experimentation. All the inhibitor treatments prior to chemoattractant stimulation in the *in vitro* experiments were performed at 37° C.

### Adoptive transfers and inflammation model

Mouse neutrophils (∼10^7^ cells) isolated from *WT* mice were labeled with either the cytosolic dye CellTracker Green CMFDA or CellTracker Red CMPTX, whereas neutrophils isolated from mice expressing GFP-NMIIA or GFP-Lifeact were used unstained. In select experiments, neutrophils were treated with inhibitors of LTB_4_ production (MK886), ROCK activity (Y27632), NMIIA activity (nBleb), Rab27a (Nexinhib20), N-SMase (GW4869), ER-golgi transfer (Exo1) or DMSO (0.2 % in volume) as vehicle control for 20-30 mins in RT. For Itgb2 visualization *in vivo*, DMSO-or inhibitor-treated neutrophils were incubated for 10 mins with Fc-block antibody (2.5 µg per ∼10^7^ cells), followed by 10 mins incubation with anti-Itgb2 antibody (M18/2 clone) conjugated to Alexa 594 (5 µg per ∼10^7^ cells), washed and adoptively transferred into the mice. Stained neutrophils (∼6-8 × 10^6^) resuspended in saline were co-injected with antibody to CD31 (clone 390; 15 µg) conjugated to either Rhodamine or eFluor450 dye through tail vein injection. For co-adaptive transfer experiments, neutrophils purified (∼6-8 × 10^6^ each) from *WT* mouse and *Alox5*^*-/-*^ were labeled with different dyes, mixed in 1:1 ratio, and adoptively transferred. The recipient mice were anesthetized by an intraperitoneal injection of a mixture of 100 mg/Kg ketamine (VET One) and 20 mg/Kg xylazine (Anased LA; VET One). ∼3-5 × 10^3^ *E. coli* BPs (∼10-15 µl) were gently injected into the hind footpad, using a syringe equipped with a 33 G needle (TSK Laboratory), without damaging the blood vessels and the tissue. In experiments involving inhibitors or ISMic, the recipient mice underwent short-term anesthesia by isoflurane inhalation (Forane; Baxter Healthcare Corp.), injected with *E. coli* BPs in the footpad and allowed to recover for 45 mins prior to the adoptive transfer of neutrophils along with the conjugated anti-CD31 antibody. In the case of *LyzM-GFP* mice, the anti-CD31 antibody was injected *IV* prior to anesthesia. One footpad was injected with saline as control, whereas the other was injected ∼1 h after with *E. coli* BPs.

### 2P-IVM and ISMic of the infected footpad

The injected footpad of anesthetized mouse was placed in a custom-designed holder to ensure stability, with the entire stage heated and maintained at 37° C throughout the imaging period. Imaging was performed by using an inverted laser-scanning two-photon microscope (MPE-RS, Olympus, Center Valley, PA, USA) equipped with a tunable laser (Insight DS+, Spectra Physics, Santa Clara, CA, USA). Excitation was performed at 850 nm for the experiments with *GFP-NMIIA* neutrophils and 810 nm for all the other experiments. Emitted light was collected by an appropriate set of mirrors and filters on 3 GaAsP detectors (bandpass filters: Blue = 410–460 nm, Green = 495–540 nm, Red = 575–645 nm). Images were acquired using a 37°C heated objective: 10X air objective (NA 0.4), 30X and 40X silicone oil immersion UPLSAPO objectives (NA 1.05 and 1.25 respectively), from Olympus. Imaging was performed by continuous acquisition of 20-25 Z-stacks (3 µm step size and at a frame rate of ∼6-10 secs between frames) for a total period of 15-20 minutes for individual field of view (hereafter called FOV). For ISMic, we used both 30x and 40x objectives, with a digital zoom of 2-4x, acquiring 8-10 Z-stacks with 2 µm step size and at a frame rate of ∼6-10 secs between frames for a total period of 5-10 minutes for individual FOV. Images were acquired using the Olympus Fluoview software and processed for further analysis using Fiji, Imaris (Bitplane) and MATLAB (Math Works). For representation in Movies S5 and S6 as well as Figures 2D, 3D and 3E, ISMic images were further 3D deconvolved using AutoQuant X3 (Media Cybernetics) involving a blind adaptive method with theoretical point spread function and five iterations.

### Analysis of neutrophil recruitment kinetics in the infected footpad

The maximal projection of Z-stacks shown in Figures 1B were acquired by 2P-IVM using a 10X air objective at fixed time points after inflammation. Maximal projections were used to quantify the neutrophils in each FOV using the Imaris spot detection function at each time point. A single FOV using the 10X air objective was sufficient to capture a large area with neutrophil recruitment response, which was followed over multiple time points. The numbers of neutrophils were normalized and represented as fold change with respect to 1 h post inflammation. The number of neutrophils observed at 1 h post inflammation observed in the footpad between adoptively transferred *WT* or *Alox5*^*-/-*^ neutrophils was comparable across experiments.

### Analysis of intravascular neutrophil behavior

First, we corrected for the motion artifacts due to heartbeat and respiration by applying to the raw time-lapse images a customized MATLAB function based on max cross-correlation image registration algorithm (Brown, 1992). Next, we determined the outline of the blood vessels for each time frame by segmentation of the maximal projections derived from the volume rendering of the acquired Z-stacks. Neutrophil contours were identified using a snake algorithm implemented with MATLAB scripts, as previously described (Chen et al., 2016; Xu and Prince, 1998), and they were overlaid to the max intensity projection of the blood vessels. To measure neutrophil velocity, we used a manual tracking plugin in Fiji. To determine their position with respect to the blood vessels we calculated the overlap fraction (OF). For each cell at each frame, the OF was calculated as the area of the 2-D max projection of the cell overlapping with the segmented blood vessels divided by the total cell area. Neutrophils were considered inside the blood vessels if the OF exceeded 0.5. Neutrophils were classified as showing free-flowing, rolling, arrest or extravasation behavior according to the following criteria: 1) free-flowing, if the velocity was higher than 40 µm/s and the OF > 0.5; 2) rolling, if the velocity was between 2 and 40 µm/s (Sundd et al., 2011) and the OF > 0.5; 3) arrest, if the velocity was lower than 0.15 µm/s and remained at the same place for 30 secs (Miyabe et al., 2017) or more with the OF > 0.5; and 4) extravasation, if the OF dropped below <0.5. The neutrophil behaviors were scored and represented independently for each cell in the field of view (FOV). As a neutrophil can show one or more behavior during the course of acquisition, each behavior was independently scored and therefore the % cells that display rolling, arrest and extravasation will not add up to 100% by this analysis. Multiple FOVs were imaged in each experiment and typically data sets between 1-2 h post inflammation were considered for analysis, with the data normalized and presented as % cells across FOVs analyzed for rolling, arrest and extravasation behavior for each experiment in each condition. For the experiments involving Y27632, nBleb and MK886 treatments, the % of cells that arrested and extravasated were scored manually from multiple FOVs in each ISMic imaging experiment.

### Analysis of peripheral distribution index and EVs

We acquired Z-stacks (typically 8-12 stacks) at a frame rate of ∼5-8 secs. Only neutrophils that displayed arrest behavior were considered for the time periods - 0, 30, 60 and 90 secs of arrest within the blood vessel. In every frame, we identified the sections corresponding to a given cell and determined the cortical redistribution of either GFP-NMIIA or Itgb2 by assigning a value of 0 for a centrally distributed fluorescent signal within the cell, and 1 for a cortically distributed fluorescent signal at the cell periphery. For each cell, we averaged the scores determined in each section of the stack to the total number of stacks analyzed across the time period and the index for each cell analyzed was determined. The diameter of EVs were measured using Imaris software. The number of EVs detected across multiple FOVs were normalized to the number of detected neutrophils in the same, for individual experiment per condition.

### In vitro adhesion assay and the analysis of actomyosin polarity

Neutrophils (2 × 10^5^ cells) were plated in 8-well chambers with a bottom coverglass coated with 100 µg/ml of fibrinogen (Lab-Tek; 1 NA). All the experiments were performed in RPMI medium and at 37° C. Cells were pretreated with vehicle (DMSO; 0.2% in volume) or specified inhibitors for 20 mins before stimulation with chemoattractants. Post stimulation, cells were gently washed with RPMI medium, fixed with 4% PFA and stained with DAPI (10 ng/ml) and rhodamine phalloidin (0.2 Units). For the analysis of cortical distribution of NMIIA, cells were permeabilized with a buffer (0.5% Saponin and 0.5% Triton X 100, both from Sigma Aldrich) for 5 mins at 37° C, followed by staining with anti-NMIIA antibody (in 1% FBS containing PBS) overnight at 4° C and thereafter with a secondary antibody for 2 h in RT. Images were acquired by using an 880 Airyscan microscope equipped with GaAsP detectors and using either 20X (Zeiss Plan Apochromat 20X/ NA 0.8, air) or 40X (Zeiss Plan Apochromat 40X/ NA 1.4, oil) lenses and acquired using the ZEN software (Carl Zeiss). The number of cells per FOV were estimated by measuring the DAPI signal using the Imaris spot detection method. Typically, more than a hundred cells per condition were analyzed from multiple FOVs. For the kinetic analysis of adhesion response reported in Figure 4A, the number cells from FOVs were averaged for each time point and represented as fold change with respect to unstimulated control. For the other experiments, results were normalized and represented as % of the vehicle treated controls. Rhodamine phalloidin and NMIIA staining was used to manually quantify the % cells in each FOV exhibiting polarized F-actin and cortical NMIIA distribution and averaged for each condition in an experiment. Typically, around 100 cells or more per condition were analyzed from multiple FOVs in each experiment to determine actomyosin polarity in PMNs. For experiments involving *GFP-NMIIA* neutrophils, cells were fixed as above but only stained with DAPI and Rhodamine Phalloidin and analyzed as mentioned above.

### Analysis of ITGB2 vesicle trafficking in vitro

For live cell imaging, Alexa 488-conjugated anti-ITGB2 antibody (αITGB2, CTB104 clone, 2 µg) and CellMask deep red (100 nM) were added to vehicle-or inhibitor-treated PMNs suspended in RPMI medium and plated on fibrinogen-coated surface, before stimulation with 100 nM fNLFNYK and thereafter imaging for ∼10 mins post-stimulation, within an incubation chamber set at 37° C. Images were acquired in time-lapse by using an 880 Airyscan microscope using the 63X (Zeiss Plan Apochromat 63X/ NA - 1.4, oil) heated lens (37° C). For the analysis of ITGB2 internalization, Alexa Fluor 488 conjugated αITGB2 (0.1 µg/ml) was added to vehicle or inhibitor pre-treated PMNs suspended in RPMI medium, before stimulating with fNLFNYK at 37° C. After 10 mins of stimulation, cells were fixed and incubated overnight at 4°C with an anti-AlexaFluor 488 functional antibody (0.5 µg per well) to quench the αITGB2 signals on the PM. Samples were counter-stained with DAPI and Phalloidin. Internalized ITGB2 was detected using either epifluorescence (Revolve FL, FJSD1000 from ECHO Laboratory) or confocal (Carl Zeiss 880 Airyscan microscope) microscopes equipped with 40X or 63X objectives. The Revolve FL epifluorescence microscope was equipped with CMOS camera and 40x (LUCPlan FL N) Olympus objective lens. The number of internalized fluorescent vesicles were determined using the Imaris spot detection method. Typically, more than a hundred cells per condition were analyzed from multiple FOVs. Results were normalized and represented as % or fold change with respect to the fluorescent vesicles detected in the control or vehicle condition. For the ITGB2 recycling experiments, vehicle-or inhibitor-treated PMNs were stimulated in the presence of Alexa Flour 488 conjugated anti-ITGB2 antibody for 15 mins. Cells were gently washed with RPMI and either fixed or incubated for an additional 45 min in RPMI containing fNLFNYK (with or without inhibitor) and anti-Alexa488 functional antibody to allow recycling of internalized ITGB2. Cells were then fixed and the αITGB2 signals on the PM were further quenched as described above. The number of internalized fluorescent vesicles were determined using the Imaris spot detection method as described above. Results are presented as the % difference in the detected fluorescent vesicles to that of 15 mins time point.

### Quantification, statistical analysis and representation

Microsoft Excel was used for calculations, the results were plotted and analyzed using Prism (GraphPad Software, Inc.). Statistical tests for each graph and the size of the samples are described in the respective figure legend. In the graphs, *p* value less than 0.05, 0.01, 0.001 and 0.0001 are represented with *, **, *** and **** respectively. For clarity, results from the IVM experiments are presented as scatter plots with bar graphs, and *in vitro* neutrophil stimulation experiments as column bar graphs, respectively.

### Online Supplementary material

Figures S1, S2 and S3 support the Figs. 1-2, 4 and 5 respectively. Figure S2K summarizes our observations on ITGB2 dynamics in PMNs. Figure S4 presents a model based on our conclusions. Supplementary movies were generated using Adobe Photoshop software to assemble individual movies into one and for labeling/annotation purposes. Movies S1-S9 have been discussed in the results section. Legends for figures and movies are provided.

## ACKNOWLEDGEMENTS

This research was supported by the Intramural Research Program of the National Institutes of Health, National Cancer Institute and Center for Cancer Research. We thank Drs. Dorian McGavern (NINDS) and Robert S. Adelstein (NHLBI), for providing us the *LyzM-GFP* and *GFP-NMIIA* mouse lines respectively. We thank the animal facilities of NCI and Dr. Wang-Ting Hsieh, FNLCR, for maintaining and genotyping the animals respectively. We also thank Dr. Michael Kruhlak, Langston Lim and Dr. Andy Tran at the CCR confocal core, for their help with confocal microscopy.

## AUTHOR CONTRIBUTIONS

Conceptualization - B.C.S, R.W and C.A.P; Methodology – B.C.S, N.M, W.W, D.G, R.W and C.A.P; Investigation – B.C.S, N.M and D.G; Analysis – B.C.S, D.C and R.W; Writing, original draft – B.C.S, R.W and C.A.P;

