## Supplementary Figure And Movie Legends for "The LTB_4_-BLT1 axis regulates actomyosin and β2 integrin dynamics during neutrophil extravasation"

**SUPPLEMENTARY FIGURE LEGENDS**

**Figure S1.**

**A)** Set up of the mouse footpad inflammation model. Upper Panel – Diagram of the 2P-IVM procedure in the *LyzM-GFP* mouse footpad. Lower panels – Mice were injected with either saline (left panel) or *E. coli* BPs (right panel) in the foot pad and imaged ~40-60 mins post-treatment by 2P-IVM. Maximum projection of Z-stacks is presented. Vasculature (blue) and neutrophils (green). Pink arrowheads mark extravasating neutrophils (right panel). Scale bar equals 40 µm. The experiments were performed in N=2 mice. See Movie S1.

**(B-D)** Neutrophils purified from *WT* mice were stained with CellTracker Green CMFDA, treated with either 40 µM Y27632 or 10 µM nBleb or the vehicle (DMSO) for ~30 mins, adoptively transferred into *Alox5*$^{-/-}$ mice with inflamed footpad and imaged by 2P-IVM. **B**-Maximum intensity projections of Z-stacks are presented. Magenta arrowheads highlight neutrophils (green) extravasating from the blood vessels (blue). Scale bar equals 20 µm. **C-D** Quantification of the % of neutrophils that exhibited arrest **(C)** or extravasation **(D)** during the late phase of response, typically 45-90 mins after neutrophil transfer to the recipient mice. Data are plotted as Means ± SEM from N=3 mice for each condition. No significant difference was observed upon analysis with unpaired *t* test with Welch's correction. Unpaired *t* test with Welch's correction was used to determine statistical significance.

**(E)** Neutrophils purified from *GFP-NMIIA* mice were treated with either the vehicle (DMSO; top row) or 5 µM MK886 (bottom row), injected into *Alox5*$^{-/-}$ mice with inflamed footpad, and imaged in time-lapse modality by ISMic. The optical sections were used to quantify the redistribution index of GFP-NMIIA. Scale bar equals 3 µm. For each time point, each section in Z axis was given a score of 0 or 1 if the GFP-NMIIA was enriched at the center or cortex of an arrested neutrophil manually. An index was generated by averaging the score to the number of stacks analyzed for each neutrophil that displayed arrest behavior during ISMic analysis. For further details please refer the Methods section.

**Figure S2.**

**(A)** PMNs were pre-treated with the vehicle or 2 µM MK886 or 20 µM LY223982 for 20 mins and stimulated with 25 nM fNLFNYK or 250 ng/ml C5a or 250 ng/ml CXCL8 for 15 mins. Adhesion was calculated as described in the Methods section and presented as % to the respective vehicle controls for each chemoattractant. Data are plotted as Means ± SEM from N=4 independent experiments. Two-way ANOVA analysis using Dunnette's multiple comparison test was used to determine statistical significance.

**(B-C)** Neutrophils from *WT* **(B & C)**, *Blt1*$^{-/-}$ **(B)** and *GFP-NMIIA* **(C)** mice were stimulated with 25 nM WKYMVm for 15 mins on a fibrinogen-coated surface. Adhesion was calculated as described in the Methods section and presented as % to the *WT* controls. Data are plotted as Means ± SEM from N=3 independent experiments in each panel. Unpaired *t* test with Welch's correction was used to determine statistical significance and no significant difference was observed between conditions in **(C)**. mNeu label stands for data from mouse primary neutrophils.

**(D-F)** *GFP-NMIIA* neutrophils were pre-treated with the vehicle or 2 µM MK886 or 20 µM LY223982 for 20 mins, stimulated with 25 nM WKYMVm for 15 mins, washed, fixed and stained with rhodamine phalloidin. **D-** Adhesion was calculated as described in the Methods section and presented as % of vehicle treated controls. **E-F** Quantification of the % of neutrophils exhibiting cortical NMIIA **(E)** and polarized F-actin **(F)** in response to above mentioned treatments. Data are plotted as Means ± SEM from N=3 independent experiments. One-way ANOVA analysis using Dunnette's multiple comparison test was used to determine statistical significance. mNeu label stands for data from mouse primary neutrophils.

**(G-I)** PMNs were pre-treated with either the vehicle or 20 µM LY223982 for 20 mins. They were incubated with an Alexa488-conjugated antibody against human ITGB2 ($\alpha$ITGB2, CTB104 clone; green) in combination with CellMask Deep Red (magenta) before stimulation with 100 nM fNLFNYK for 10 mins on a fibrinogen-coated surface and acquired in 3D using time-lapse using confocal microscopy. **G-** The trajectory of the internalized vesicles was determined as described in the Methods section. Tracks of individual ITGB2-containing vesicles between 5-7 mins post stimulation are over-layed on the maximum intensity projections derived from Z-stacks. Scale bar equals 5 µm. See

Movie S7. **H-I** Quantification of the number **(H)** and distance **(I)** travelled by ITGB2-containing vesicles for each condition. Data are plotted for individual cells with n=16 cells in vehicle and n=18 cells in LY223982 treatment from N=3 independent experiments. Unpaired *t* test with Welch's correction was used to determine statistical significance.

**(J-K)** PMNs were pre-treated for 20 min with vehicle (DMSO) or the indicated inhibitors of clathrin and dynamin (5 μM PitStop2 or 100 μM MDC or 50 μM Dynasore or 2.5 μM MitMAB), stimulated for 10 min with 25 nM fNLFNYK, fixed and imaged by confocal microscopy. **J**- Representative confocal images of PMNs with internalized ITGB2 under indicated conditions is presented. White dashed lines indicate cell boundary and scale bar equals 5 μm. **K**- The extent of ITGB2 internalization was determined as described in the Method section and the data are plotted as % change with respect to the vehicle. Data is represented as mean ± SEM from N=3 independent experiments. One-way ANOVA analysis using Dunnette's multiple comparison test was used to determine statistical significance.

**(L)** A model of ITGB2 trafficking regulated by the $LTB_4$-BLT1 axis in PMNs. The figure depicts the progression of PMN adhesion on a fibrinogen-coated surface. In control condition, ITGB2 clusters and migrates to the bottom of the cell to form a ring-like structure, which then disassembles as the PMN begins to migrate. However, upon blockade of $LTB_4$ sensing via BLT1, clusters of ITGB2 rapidly internalize and fail to assemble into a ring-like structure at the bottom of the PMN, resulting possibly in the deadhesion of PMN with time.

**Figure S3.**

**(A-C) A**- A diagram of the 2P-IVM procedure in *Alox5*$^{-/-}$ mouse with inflamed footpad, where neutrophils from *WT* (green) that were treated as indicated in Fig. 5A for 30 mins before they were co-injected with untreated *Alox5*$^{-/-}$ neutrophils into the recipient. **B**- An illustration of the possible autocrine/paracrine $LTB_4$ signaling that occurs between *WT* and *Alox5*$^{-/-}$ neutrophils during extravasation response when co-injected into the recipient mice with inflamed footpad, and how this mechanism can be blocked using the $LTB_4$ inhibitor MK886. **C**- An illustration of the possible EV-based autocrine/paracrine $LTB_4$ signaling in the conditions mentioned above that occurs in the inflamed vasculature, and how this mechanism can be blocked using inhibitors to N-SMase and Rab27. See Figs. 5A and 5B as well as Movies S8 and S9.

**(D-G)** PMNs were pretreated for 20 mins with vehicle (DMSO) or 10 μM GW4869 or 1 μM Nexinhib20 or 10 μM Exo1 and stimulated in the absence or presence of $\alpha$ITGB2 for 15 min with 25 nM fNLFNYK, fixed, stained and imaged by confocal microscopy. Quantification of the % of cells with cortical NMIIA distribution **(D)** and polarized F-actin **(E)** is presented. The extent of internalized ITGB2 **(F)** and cell adhesion **(G)** observed under the above-mentioned conditions were measured and represented as fold change and % with respect to vehicle control, respectively. Data are plotted as mean ± SEM from N=3 independent experiments. One-way ANOVA analysis using Dunnette's multiple comparison test was used to determine statistical significance.

**Figure S4.**

Model depicting the role of the $LTB_4$-BLT1 axis in promoting neutrophil arrest and extravasation *in vivo*. Under control conditions, neutrophils respond to inflammatory cues by releasing EVs that relay $LTB_4$ signals in an autocrine/paracrine manner. The $LTB_4$-BLT1 axis triggers the redistribution of NMIIA and Itgb2 for sustained arrest in the inflamed vessel followed by extravasation into the interstitium. These mechanisms are impaired in *Alox5*$^{-/-}$ or *Blt1*$^{-/-}$ neutrophils, resulting in their reduced arrest and extravasation in response to inflammation. The specific mechanisms regulating the release of $LTB_4$ from EVs and the local action of $LTB_4$ in the inflamed vessels have yet to be determined.

### SUPPLEMENTARY MOVIE LEGENDS

**Movie S1.** 2P-IVM of the *LyzM-GFP* mouse footpad at ~1 h post injection of saline in one footpad and *E. coli* BPs in the other. Movies are maximum intensity projection of an image stack acquired for a period of ~ 20 mins (min:sec:ff) in each footpad with a frame rate of 10 frames per second (fps). The vessels were labeled with anti-CD31 (blue). Scale bar equals 50 µm.

**Movie S2.** 2P-IVM of the footpad in *WT* mouse, that received labeled *WT* neutrophils, at ~1.5 h post injection of saline in one footpad and *E. coli* BPs in the other. Movies are maximum intensity projection of an image stack acquired for indicated time periods (min:sec:ff) in each footpad with a frame rate of 10 fps. The vessels were labeled with anti-CD31 (blue). Scale bar equals 50 µm.

**Movie S3.** 2P-IVM of the inflamed *Alox5*$^{-/-}$ mouse footpad at ~1.5 h, where labeled neutrophils (green) were introduced from *WT* or *Alox5*$^{-/-}$ or *Blt1*$^{-/-}$ mice. Movies are maximum intensity projection of an image stack acquired for a period of ~ 15-20 mins (min:sec:ff) for each condition with a frame rate of 10 fps. The vessels were labeled with anti-CD31 (blue). Scale bar equals 50 µm.

**Movie S4.** 2P-IVM of the inflamed *Blt1*$^{-/-}$ mouse footpad at ~1.5 h, where labeled neutrophils (green) were introduced from *WT* or *Alox5*$^{-/-}$ or *Blt1*$^{-/-}$ mice. Movies are maximum intensity projection of an image stack acquired for a period of ~ 15-20 mins (min:sec:ff) for each condition with a frame rate of 10 fps. The vessels were labeled with anti-CD31 (blue). Scale bar equals 50 µm.

**Movie S5.** ISMic imaging of the inflamed footpad of *Alox5*$^{-/-}$ mouse that received either vehicle or MK886 treated *NMIIA GFP* neutrophils (white), ~1 h post inflammation. Movies are single slice projection from an image stack and represent ~ 2-4 mins (min:sec:ff) of acquisition for each condition with a frame rate of 5 fps. The vessels were labeled with anti-CD31 (blue). Scale bar equals 3 µm.

**Movie S6.** ISMic imaging of the inflamed footpad of *Alox5*$^{-/-}$ mouse that received *WT* or *Alox5*$^{-/-}$ neutrophils labeled for cytoplasm (CellTracker; magenta) and Itgb2 (M18/2; white), ~1 h post inflammation. Blood vessels are labeled with anti-CD31 (blue). Movies are maximum intensity projection of an image stack acquired for a period of ~ 6-9 mins (min:sec:ff) of acquisition with a frame rate of 4 fps. Scale bar equals 5 μm.

**Movie S7. A**- Confocal imaging of PMNs stained for plasma membrane (magenta) and $\alpha$ITGB2 (white) that were pre-treated with vehicle (DMSO) or BLT1 inhibitor (LY223982) for 20 mins before stimulation with fNLFNYK and simultaneous live acquisition for 10 mins. Movies are projections of image stacks acquired in 3D, for a period of ~ 10 mins (min:sec:ff) with a frame rate of 5 fps. **B**- A view of a section from the cell bottom for the conditions mentioned in A.

**Movie S8.** 2P-IVM of the inflamed footpad at ~2 h, where labeled *WT* neutrophils (green) that were pre-treated with vehicle (DMSO) or the inhibitors MK886 or GW4689 or Nexinhib20 or Exo1 were co-injected with labeled *Alox5*$^{-/-}$ neutrophils (strawberry) into *Alox5*$^{-/-}$ mice. Movies are maximum intensity projection of an image stack acquired for a period of ~ 20-22 mins (min:sec:ff) for each condition with a frame rate of 15 fps. The vessels were labeled with anti-CD31 (blue). Scale bar equals 50 μm.

**Movie S9.** ISMic imaging of *WT* neutrophils introduced into *Alox5*$^{-/-}$ mouse with inflamed footpad ~2 h, where EVs contain CellTracker Green CMFDA (green) and Itgb2 (M18/2 clone; red). Movies are maximum intensity projection of an image stack and represent ~ 11 mins (min:sec:ff) of acquisition. The vessels were labeled with anti-CD31 (cyan). Scale bar equals 3 μm. The EVs roll along and occasionally tether to the inflamed endothelial surface facing the vascular lumen in the vicinity of extravasating neutrophils.
