## Supplementary figures and images for "The LTB_4_-BLT1 axis regulates actomyosin and β2 integrin dynamics during neutrophil extravasation"

### Supplementary Figure S1

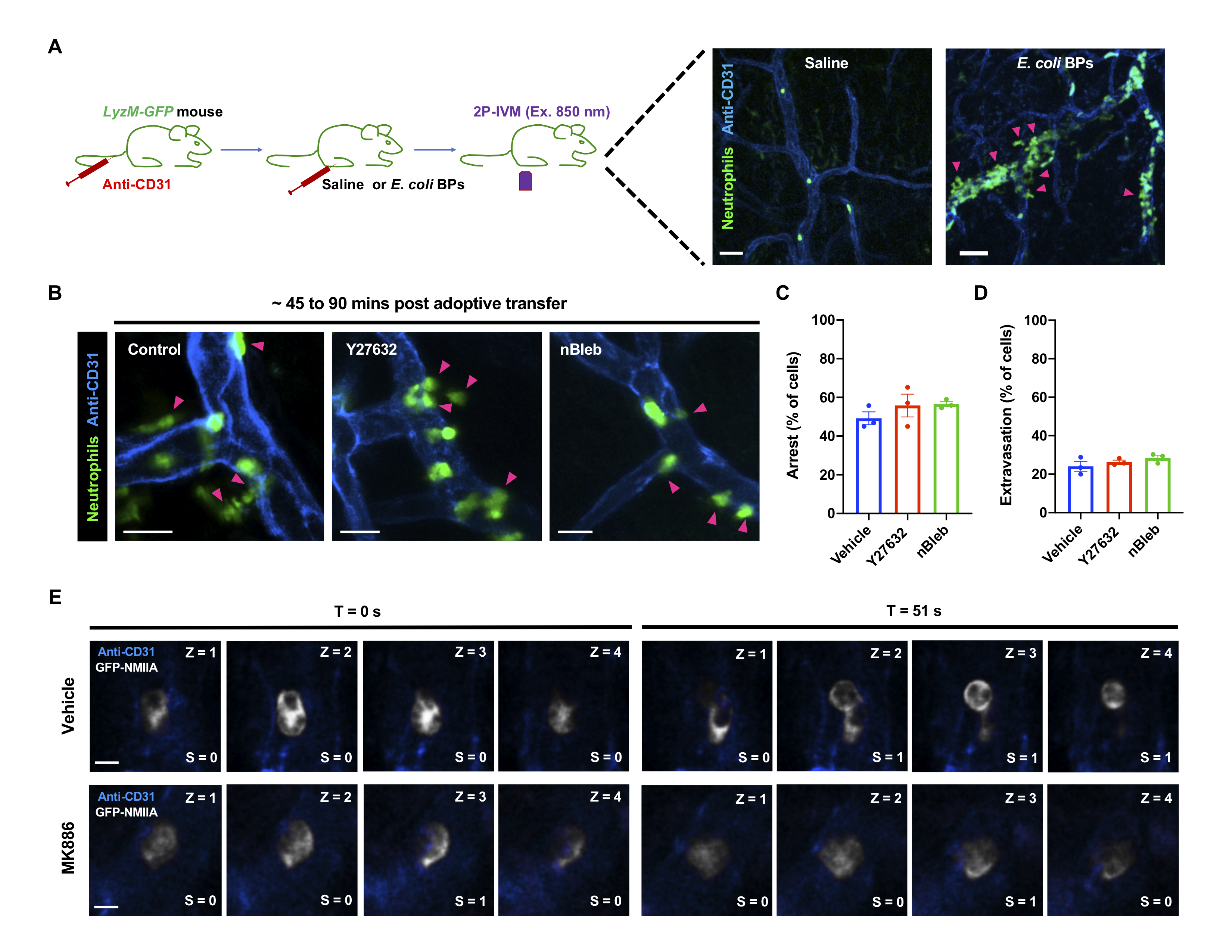

### Supplementary Figure S2

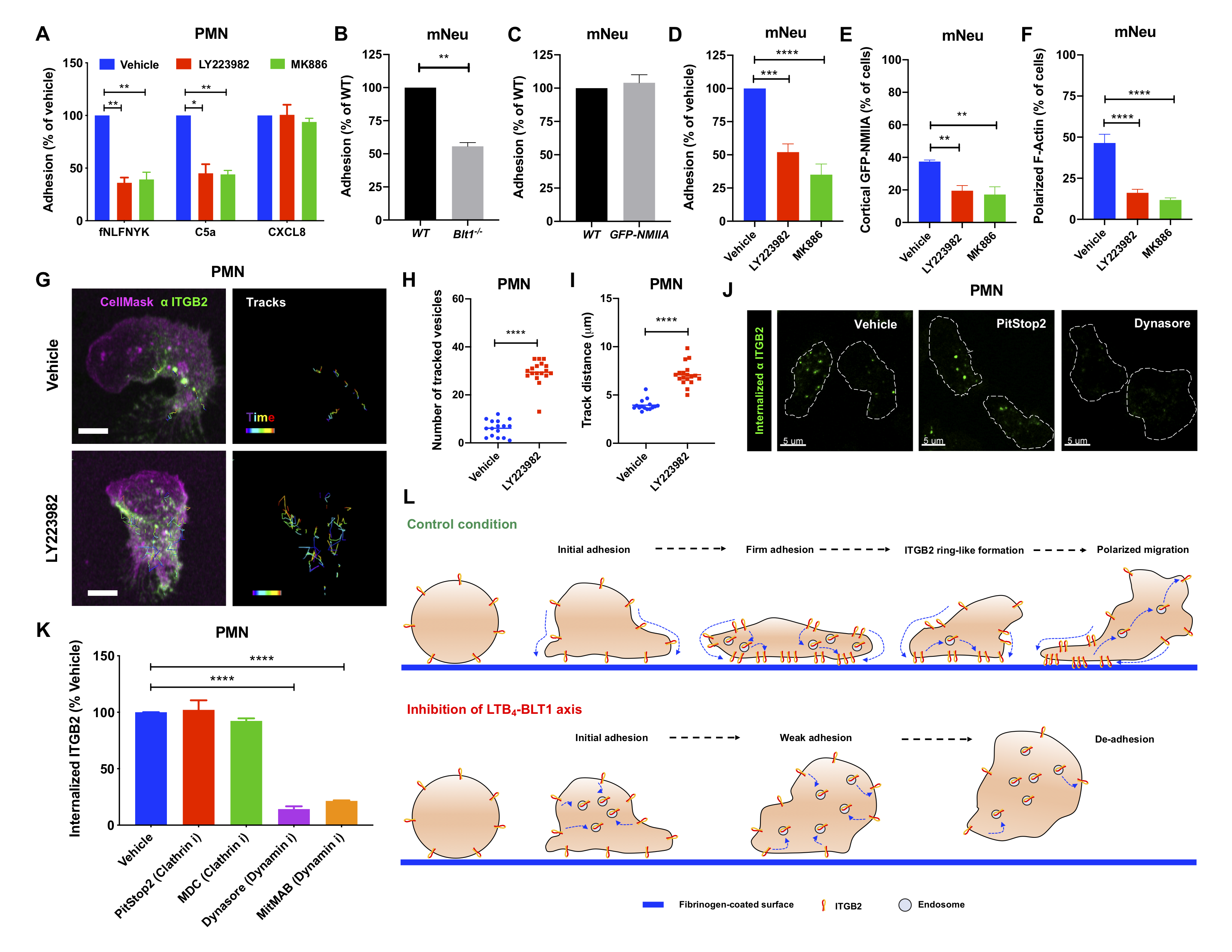

### Supplementary Figure S3

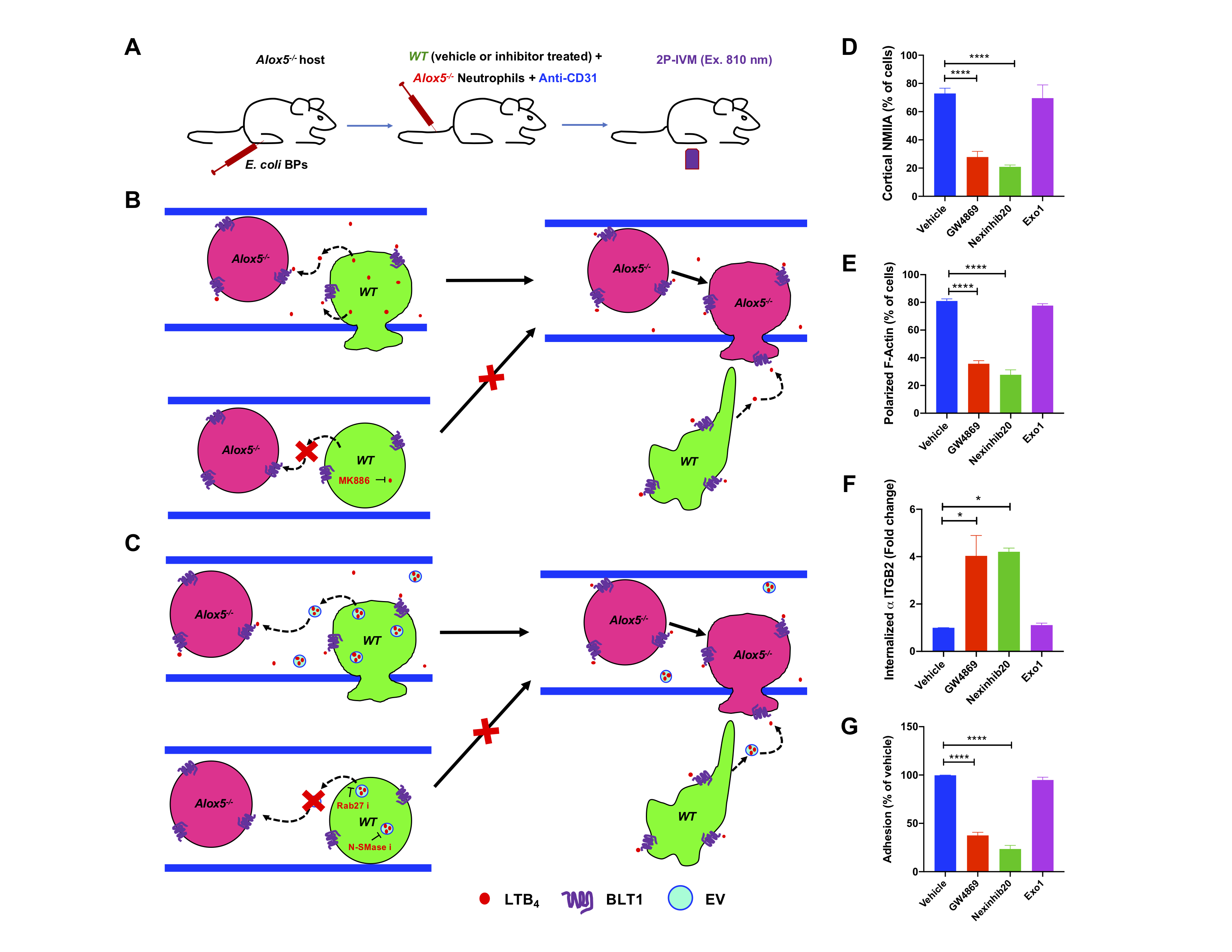

### Supplementary Figure S4

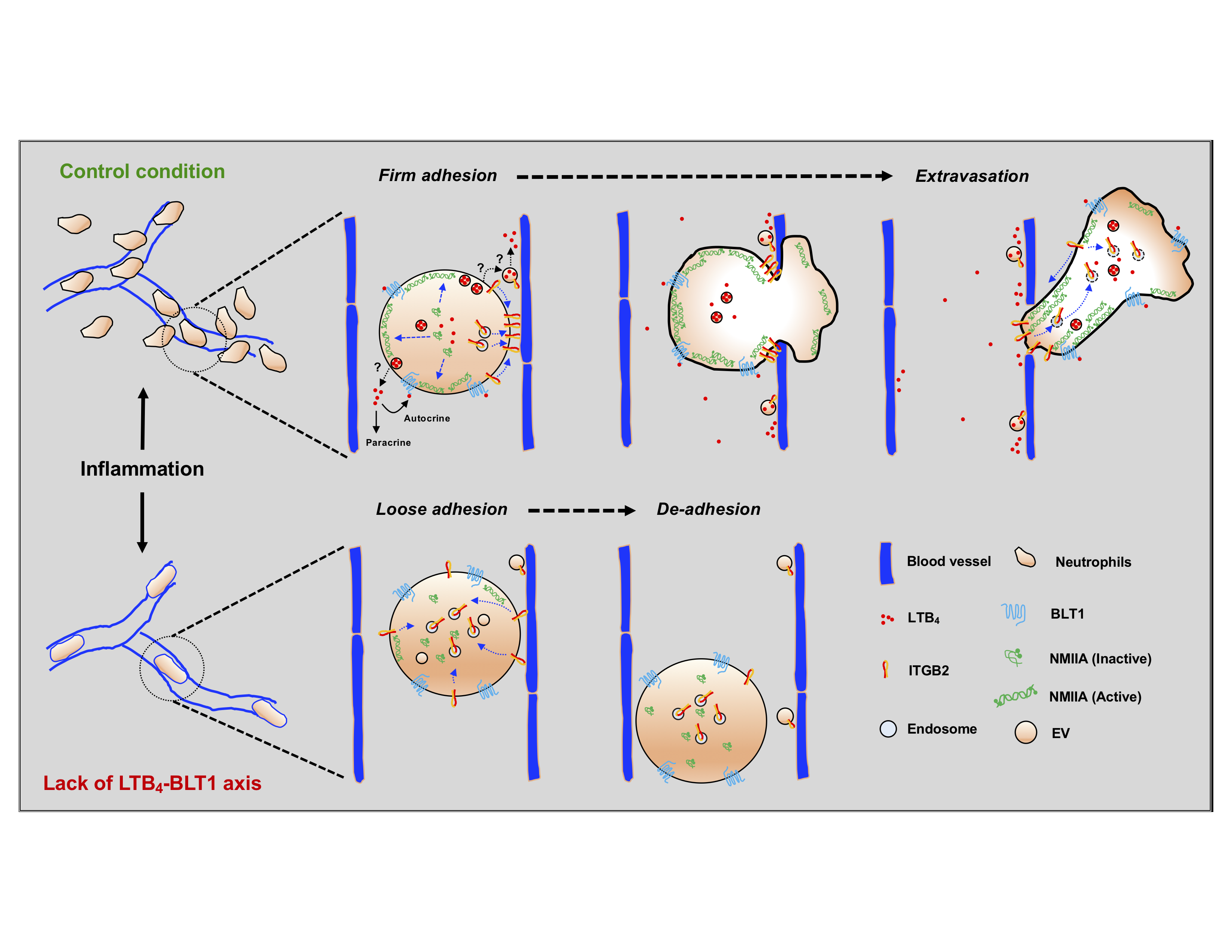
